# Molecular pathway of mitochondrial preprotein import through the TOM-TIM23 supercomplex

**DOI:** 10.1101/2023.06.21.546012

**Authors:** Xueyin Zhou, Yuqi Yang, Guopeng Wang, Shanshan Wang, Dongjie Sun, Xiaomin Ou, Yuke Lian, Long Li

## Abstract

Most mitochondrial proteins need to be imported from the cytosol. Over half of mitochondrial proteins are imported through the pre-sequence pathway that is controlled by the TOM complex in the outer membrane and the TIM23 complex in the inner membrane. It is unclear on the molecular level how proteins cross the mitochondrial double membranes through the TOM and TIM23 complexes. Here, we report the assembly of the active TOM-TIM23 supercomplex with translocating polypeptide substrates captured in the import pathway. Electron cryo-microscopy (Cryo-EM) analyses reveal that during translocation across the outer membrane, the polypeptide substrates pass through the center of the Tom40 channel while interacting with a glutamine-rich patch in the inner wall of Tom40. Structural and biochemical analyses show that the TIM23 complex contains a heterotrimer of the subunits Tim23, Tim17, and Mgr2 in the inner membrane. Tim17 and Mgr2 shield the polypeptide substrates from the lipid environment. The import pathway consists of two highly conserved residue patches of Tim17, one negatively charged patch at the entrance and one hydrophobic patch in the middle of the pathway. These data reveal an unexpected pre-sequence pathway mediated by the TOM-TIM23 supercomplex for facilitating protein import across the double membranes of mitochondria.

## Introduction

Most mitochondrial proteins are encoded in the nucleus, synthesized by ribosomes in the cytosol, and imported into mitochondrial compartments ^1, 2^. Mitochondria have evolved sophisticated import systems to facilitate protein translocation across their membranes ^3–6^. The translocase of the outer membrane (TOM) complex mediates the entrance of most proteins into mitochondria. After passing through the TOM complex, proteins are sorted and delivered to different protein translocation machinery according to their destination. Over half of mitochondrial proteins follow the pre-sequence pathway to the translocase of the inner membrane 23 (TIM23) complex ^7–9^. These protein substrates usually contain a positively charged pre-sequence at the N-terminus that directs preproteins to the matrix through the TOM complex and the TIM23 complex ^10, 11^.

The TOM complex is the primary import gate for mitochondrial proteins ^12^. The core channel-forming subunit is an integral membrane β barrel protein, Tom40. Protein substrates are translocated across the outer membrane by passing through the center of the β barrel ^13^. Tom40 form dimers and higher oligomers with Tom22, Tom20, Tom70, and small TOM subunits, including Tom5, Tom6, and Tom7. Cryo-EM structures of TOM complexes from yeast and human reveal extensive interactions among these subunits ^14–18^. In the dimeric TOM complex, two Tom22 molecules are located at the dimer interface and are involved in interactions with both Tom40 molecules. Tom5, Tom6, and Tom7 are associated with the outer wall of Tom40 at an equal ratio. Tom20 and Tom70 serve as receptors for preproteins from the cytosol ^19–22^. Tom22 can also function as a receptor for pre-sequences ^23^.

After crossing the TOM complex, protein substrates containing pre-sequences are passed to the TIM23 complex in the inner membrane. The core subunits of TIM23 are Tim23 and Tim17 which are both four-membrane-spanning proteins sharing sequence and topology similarities ^24^. While the four transmembrane segments (TMs) of either Tim23 or Tim17 alone are unlikely to complete a protein-conducting channel, Tim23 has been shown to form channels, probably by oligomerization ^25, 26^. It is generally believed that Tim23 is the major channel-forming subunit, with Tim17 providing essential structural support ^27, 28^. However, this view has recently been challenged ^29^. The other subunits of TIM23 include Tim50, Tim21, Tim44, mtHsp70, Pam16, Pam18, Mge1, Pam17, and Mgr2. The IMS domain of Tim50 recognizes pre-sequences and transports the protein substrates from the TOM complex to the TIM23 complex ^30–32^. Tim21 is suggested to regulate subunit association and activity of TIM23 ^33, 34^. Tim44 is associated with Tim23 and Tim17 at the matrix side to recruit mtHsp70 ^35–38^, the ATP-driven motor for protein translocation ^39, 40^. The activity of mtHsp70 is regulated by membrane-anchored Pam16-Pam18 and by Mge1 in the matrix ^41–47^. The most recent subunit to be identified in the TIM23 complex is Mgr2, which has two TMs and shares sequence homology with Tim17 and Tim23 ^48^. Mgr2 is a nonessential subunit that functions as a gatekeeper to regulate the lateral release of TMs from the translocase ^49, 50^.

In the pre-sequence pathway, the TOM and TIM23 complexes are orchestrated to achieve efficient protein import across mitochondrial double membranes. It is well established that TOM and TIM23 form a supercomplex during protein translocation ^51–53^. However, it is unclear how the two complexes and their subunits are organized to facilitate the threading of protein substrates through the two membranes ^54^. In particular, the composition and assembly of the core translocase unit in the TIM23 complex are obscure. To better understand the molecular details of the TOM and TIM23 complexes, particularly the import pathway during import, we assembled the TOM-TIM23 supercomplex *in vivo* with a series of polypeptide substrates. By utilizing cryo-EM, AlphFold2 modeling, site-specific photo-crosslinking, and mutagenesis techniques, we were able to visualize the import pathway of the substrates as they are translocated through the central pore of the TOM complex, and the organization of the TIM23 core that creates a cross-membrane path to facilitate protein translocation through the inner membrane.

## Results

### Architecture of the TOM-TIM23 supercomplex

The TOM-TIM23 supercomplex could be detected when the polypeptide substrates with pre- sequences are trapped in the import pathway, such as using Cytochrome b2 precursor (pB_2_) fused with dihydrofolate reductase (DHFR) as the substrate in the protein import assays of mitochondria *in vitro*^52^. pB_2_ contains a bipartite pre-sequence. The N-terminal 31 residues target the matrix and the subsequent segment (residues 32-80) contains a sorting signal to the IMS. An internal deletion of 19 residues (residues 47-65, Δ19) can disrupt the sorting signal, resulting in translocation of pB_2_ directly to the matrix ^55^. While loosely folded DHFR can be imported together with pB_2_ into mitochondria, induced folding of DHFR by methotrexate can stall protein import at the TOM complex. When the sorting signal of pB_2_ is disrupted by the Δ19 mutation in the fusion proteins and methotrexate is present, the arrested pB_2_Δ19-DHFR in mitochondria is shown to occupy the pre-sequence import pathway and trap the TOM-TIM23 supercomplex ^52^. For structural studies, we replaced DHFR within the substrate with the superfolder green fluorescent protein (sfGFP) ^56^, which stalls mitochondrial import *in vivo*. pB_2_Δ19 was truncated at residue 167, which was long enough to span the mitochondrial double membranes and could be stably trapped in both the TOM and TIM23 complexes (pB_2_167Δ19-sfGFP, Fig. 1A). The substrate was overexpressed in *Saccharomyces cerevisiae* (*S*. *cerevisiae*). The purified supercomplex contained essentially all the subunits of the TOM and TIM23 complexes as analyzed by Blue Native PAGE and mass spectrometry (Fig. 1B). Most of the subunits were evident on SDS-PAGE (Fig. S1A). The presence of the translocase core subunits, Tom40, Tim23, and Tim17, was confirmed by immunoblotting (Fig. S1B).

**Fig. 1.**
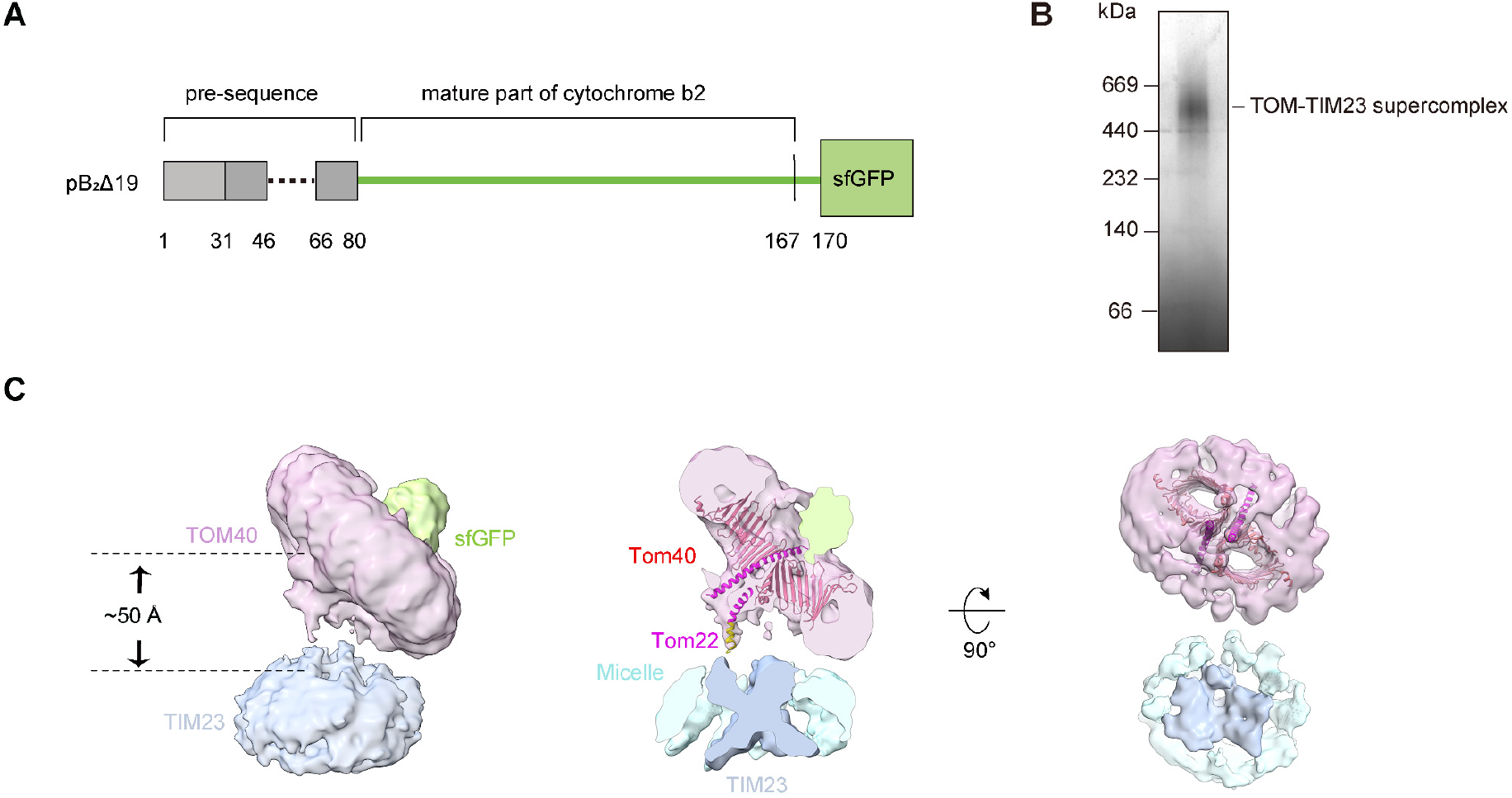
Cryo-EM structure of the TOM-TIM23 supercomplex. (**A**) Scheme of a polypeptide substrate for the assembly of the TOM-TIM23 supercomplex. The pre-sequence of cytochrome b2 precursor (pB_2_) is drawn as a bar in which the matrix targeting signal is colored light grey and the inner membrane sorting signal is colored dark grey. The mature segment of the following pB_2_ sequence is drawn as a green line. The C-terminal superfolder GFP is shown as a green box. The structural elements are labeled. The starting and ending residues of each element are marked. The dashed line indicates the Δ19 deletion in the pre-sequence. (**B**) Coomassie blue staining of the purified supercomplex analyzed by Blue Native PAGE. (**C**) Cryo-EM density of the supercomplex. Left panel, the TOM40 disc is colored pink, the TIM23 disc is colored sky blue, and GFP is colored green. The connection distance between TOM40 and TIM23 is marked. Middle panel, the cutaway sideview of the supercomplex. The TOM40 structure (PDB ID: 6UCU) is fitted in the map. The Tom40 subunit is colored red. Tom22 is colored magenta for the segment modeled in the TOM40 structure and yellow for the C terminal segment modeled by Alphafold2. The transmembrane region of TIM23 is colored sky blue and the micelle density is color light blue. Right panel, the top view of the TOM40 and TIM23 discs, showing the characteristic twin pores of TOM40 and the segmented TIM23 density.

Analysis of the supercomplex by single particle cryo-EM yielded a density map at 9.4 Å resolution (Fig. S1C-G). The supercomplex has two distinct discs of the membrane protein-detergent micelle complexes (Fig. 1C). The centers of the two discs are ∼ 50 Å apart. 3D classification revealed that the two discs are not parallel in a fixed position, but are tilted at angles of ∼30 - 50° relative to each other (Fig. S1E). The large disc has two holes in the middle that fit well with the dimeric Tom40 subunits in the TOM complex (Fig. 1C). The connection density between the large and small discs is evident, but flexible. The density could be fitted with the C-terminal helix of Tom22 which might help to connect the two discs ^53^ (Fig. 1C, S1H). The cryo-EM density of the small disc does not resolve the individual transmembrane segments (TM) of the TIM23 complex. However, the strong core density in the center of the disc has a characteristic X shape that spans the membrane (Figs. 1C, S1I). Shown as a cylinder-shaped density, the C-terminal sfGFP molecule of the polypeptide substrate is located on the cytosolic side of the TOM complex, fulfilling its function as the translocation blocker. A weak thread of density extends from sfGFP into one Tom40 channel (Fig. 1C). Thus, only one of the two Tom40 subunits in the structure is actively engaged in protein import.

### TOM complex engaged with the polypeptide substrate

The flexibility of the supercomplex prevented the further improvement of the map resolution. In an attempt to restrain the substrate in the supercomplex, a DHFR molecule was introduced in the middle of the polypeptide substrate between residues 125 and 126, a position that would allow DHFR to be translocated and folded on the matrix side of TIM23 (Fig. 2A). With this substrate (pB_2_167Δ19- DHFR-sfGFP), TOM and TIM23 would be sandwiched between sfGFP and DHFR in the supercomplex. Indeed, the cryo-EM map of this supercomplex shows that while the overall architecture was similar to the supercomplex with the substrate pB_2_167Δ19-sfGFP described above, an extra density from DHFR was visible on the matrix side of the TIM23 disc (Fig. 2B). The cryo-EM structure of this supercomplex was determined at 10.0 Å resolution (Fig. S2A-D); however, the subsequent calculation focusing on the TOM disc yielded a structure at 4.1 Å resolution (Fig. S2E-H). In the structure, it is well resolved the β barrels of the core Tom40 channels and the associated α helical Tom subunits, including Tom22, Tom5, Tom6, and Tom7 (Figs. 2C, D, S3). The two Tom40 subunits are almost identical in the structure. Similar to the structure described above, only one Tom40 channel has sfGFP trapped on the cytosolic side. The β strands of sfGFP are not resolved in the density map, suggesting that sfGFP may not be in a fixed orientation.

**Fig. 2.**
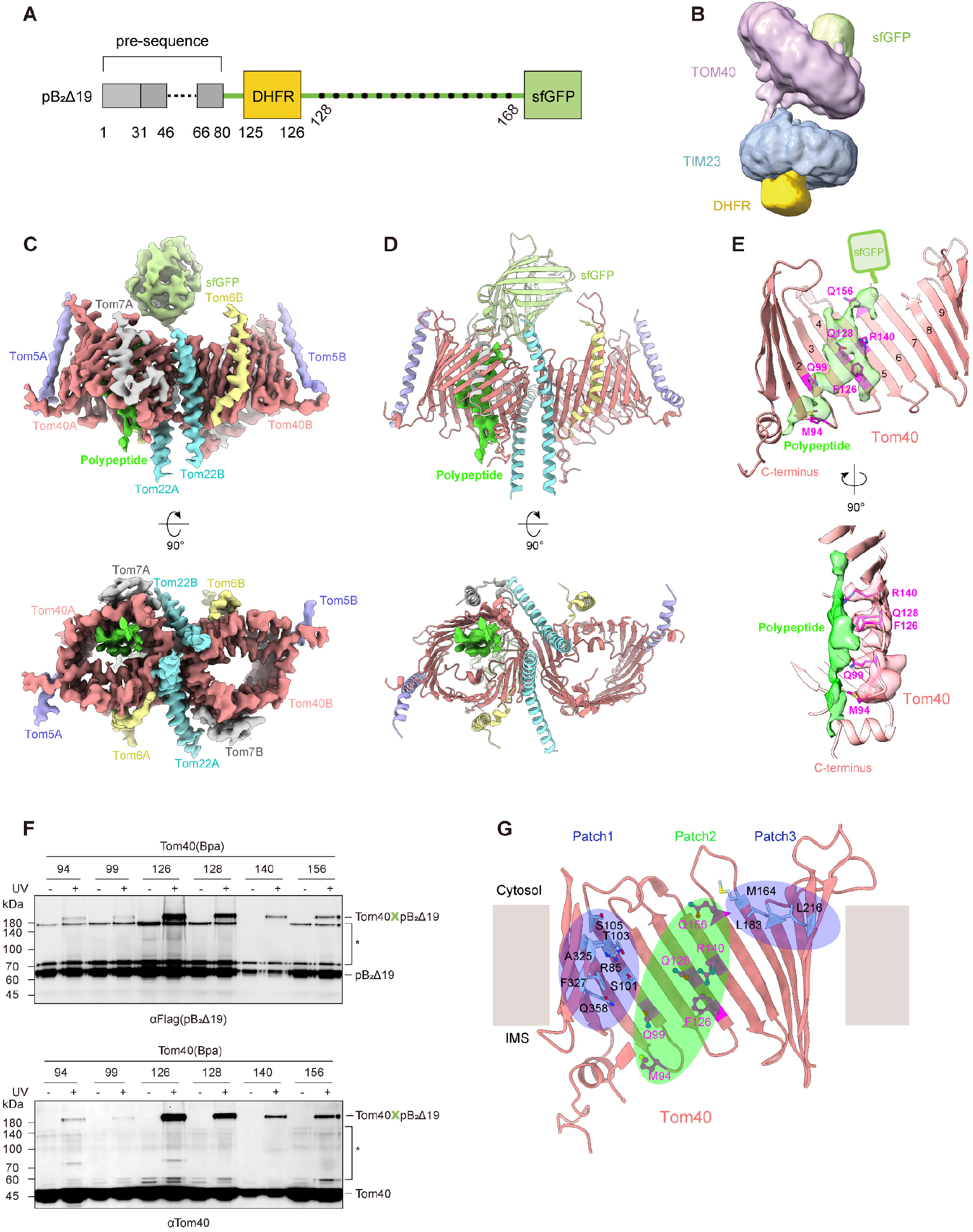
Structure of the substrate-engaged TOM40 complex. (**A**) Scheme of the DHFR containing polypeptide substrate (pB_2_167Δ19-DHFR-sfGFP) for the assembly of the TOM-TIM23 supercomplex. DHFR is drawn as a yellow box. The insertion position of DHFR in the substrate is marked. The mutation positions for photo-crosslinking are indicated by black dots. The starting and end positions are labeled. (**B**) Cryo-EM density map of the supercomplex assembled with pB_2_167Δ19- DHFR-sfGFP. DHFR on the matrix side of the structure is colored yellow. (**C**) Cryo-EM density map of the substrate-engaged TOM40 complex, showing the side view (top panel) and the top view (bottom panel). Individual subunits are labeled and colored, Tom40 in salmon, Tom22 in cyan, Tom5 in purple, Tom6 in light yellow, Tom7 in grey, GFP in light green, and the polypeptide substrate in green. To highlight the polypeptide substrate, the display threshold for the substrate density is lower than that for other subunits. (**D**) Ribbon diagrams of the substrate-engaged TOM40 complex. The coloring of the subunits is the same as in **C**. The polypeptide substrate passing one of the Tom40 channels is shown as density. (**E**) Polypeptide substrate passing the inner wall of Tom40, shown in two views that differ by 90 ° rotation. The polypeptide is shown as the density in green. The residues that interact with the substrate are shown as magenta sticks. The position of GFP is indicated by a green box. The β strand numberings of Tom40 are marked in black. The C-terminus of Tom40 is labeled. In the lower panel, the density of the substrate interacting residues is shown as a transparent salmon surface. (**F**) Photo-crosslinking of the Tom40 residues to the polypeptide substrate pB_2_167Δ19-DHFR-sfGFP (pB_2_Δ19) in the supercomplex assembled in yeast strains yLS510/ pLSC1-X. The crosslinking bands are confirmed by immunoblotting of the Flag tag (substrate) and Tom40. Non-specific bands are indicated by *. (**G**) Three patches of the substrate-interacting residues in Tom40. As revealed by a previous photo-crosslinking study ^13^, the residues that interact with the pre- sequence and carrier proteins are clustered in patches 1 and 3. These residues are shown as cyan sticks and are shaded by blue ovals. The substrate interacting residues from the cryo-EM structure are colored magenta and shaded by a green oval.

**Fig. 3.**
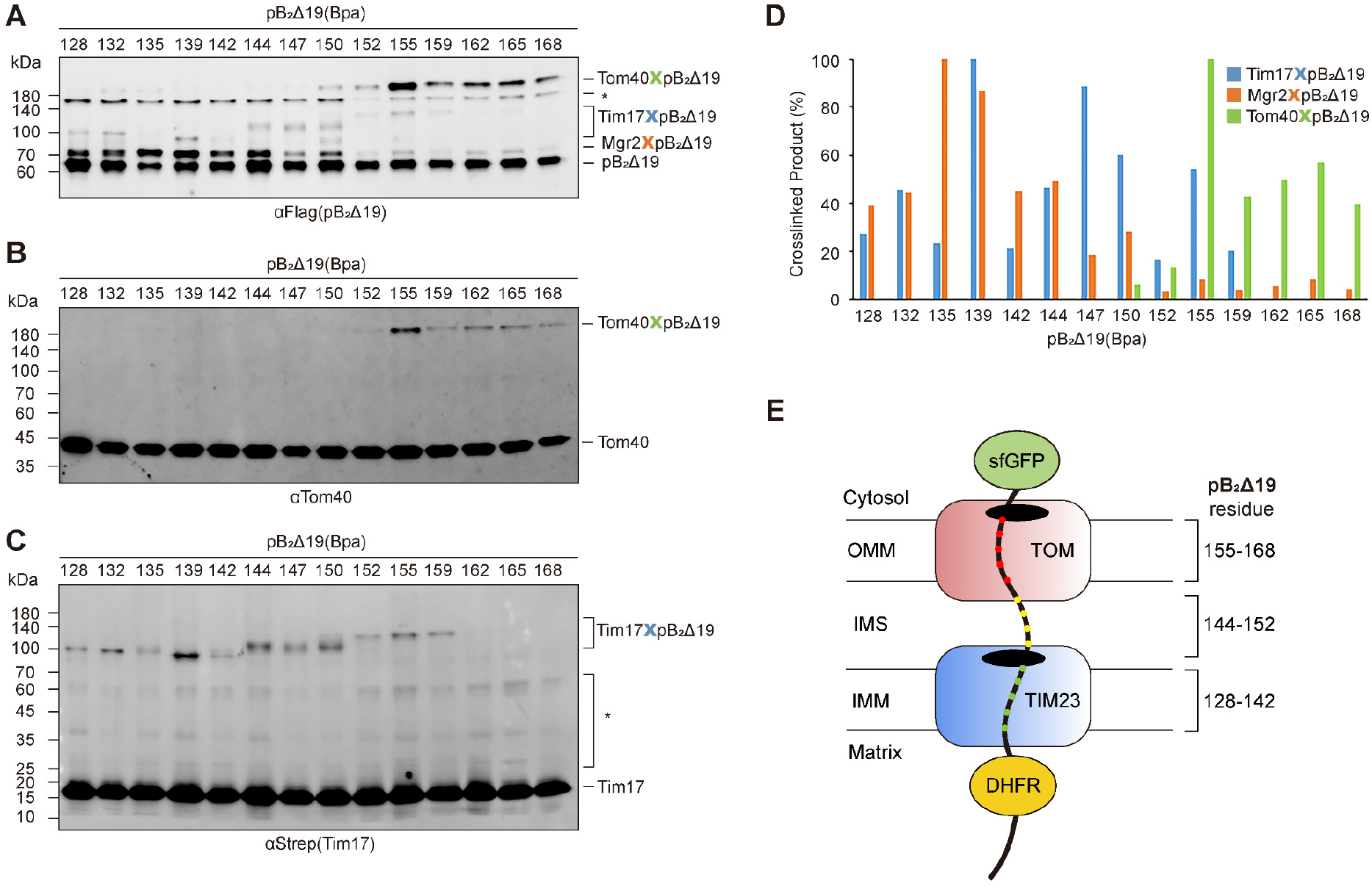
Mapping of the polypeptide substrate in the supercomplex by photo-crosslinking. (**A**) Photo-crosslinking of the substrate to the subunits in the TOM40 and TIM23 complexes. The substrate pB_2_167Δ19-DHFR-sfGFP (pB_2_Δ19) was mutated at the indicated residue positions for Bpa incorporation. The supercomplexes assembled with different pB_2_Δ19 mutants in yeast strains yLS210/pLSB4-X were purified and subjected to UV irradiation. Immunoblotting detects the Flag tag of the substrate. The crosslinking bands with different subunits are marked. (**B**) Same protein samples as in **A**, with immunoblotting of the Tom40 subunit. (**C**) Same protein samples as in **A**, with immunoblotting of the Strep tag fused to the Tim17 subunit. (**D**) Quantitation of the crosslinking efficiency based on the band intensity in **A**. The strongest crosslinking bands of each subunit over the substrate bands in the same lane were set to 100%. Band intensity was quantitated by using ImageJ. (**E**) Scheme of substrate mapping in the TOM-TIM23 supercomplex. The residue ranges of the substrate in the outer membrane, IMS, and inner membrane are marked according to the photo- crosslinking results.

A trace of density from the polypeptide substrate extends from sfGFP into one of the Tom40 channels (Fig. 2C, D). The density is evident within the channel, but not well resolved enough to build a polypeptide model, indicating that the substrate is extended and flexible in the translocation pathway (Fig. 2E). The substrate runs along a hydrophilic patch of the Tom40 inner wall that consists of β2-6. Specifically, multiple polar residues, particularly glutamine, within the patch make direct interactions with the substrate (residues Q99, Q128, R140, and Q156) (Fig. 2E). Glutamine and arginine can form extensive hydrogen bonds with a variety of residues via their long polar side chains. In addition, two hydrophobic residues, F126 and M94, are located along the pathway, probably providing hydrophobic interactions with the hydrophobic residues in the extended polypeptide substrate. The pathway was confirmed by photo-crosslinking between the residues and the polypeptide substrate (Fig. 2F).

Interestingly, this patch of the residues along the import pathway does not overlap with, but are instead flanked by, the polar and hydrophobic residues previously identified to interact with pre- sequences and hydrophobic carrier proteins ^13^ (Fig. 2G). Thus, Tom40 appears to employ different patches of residues to coordinate the translocation of different types of polypeptide substrates. After passing the central pore, the polypeptide substrate runs toward the C-terminus of the Tom40 subunit (Fig. 2E), consistent with the observation that deletion of the C-terminus of yeast Tom40 reduces import efficiency ^14^.

### Mapping of the polypeptide substrate in the supercomplex

Due to the resolution limit of the cryo-EM maps and the intrinsic flexibility of the supercomplex, the amino acid sequence of the polypeptide substrate could not be accurately assigned in the structure. We turned to the photo-crosslinking assays to map the residues of the polypeptide substrate in the import pathway. The photoreactive amino acid, 4-Benzoyl-L-phenylalanine (Bpa), was introduced into the polypeptide to probe the subunits of the TOM and TIM23 complexes in the immediate vicinity along the import pathway. pB_2_167Δ19-DHFR-sfGFP was systematically mutated to yield a series of mutants in which one in every two to four residues of the polypeptide segment between sfGFP and DHFR was mutated to the amber codon for Bpa incorporation (Fig. 2A). The supercomplexes were assembled with the Bpa containing substrates. After purification, the supercomplexes were subjected to UV irradiation. Strong crosslinking bands to Tom40 were observed for the C-terminal polypeptide segment from residues 155 to 168 (Figs. 3A, B, S4A). This segment would span a linear distance of ∼39 Å in an extended conformation, corresponding well with the thickness of the Tom40 barrel (∼ 38 Å).

In the N-terminal segment of the polypeptide, while the Tom40 crosslinked bands disappeared, prominent crosslinked bands around 100 kDa began to emerge, from residues 128-159 (Figs. 3A, S4B). These bands turned out to be Tim17 crosslinking products (Fig. 3C). Interestingly, residues 155 and 159 could be crosslinked to both Tom40 and Tim17, indicating that parts of Tim17 extend into the IMS and toward Tom40 (Fig. 3B-D). In the IMS, Tim17 has a C-terminal tail of 19 amino acids (residues 140-158) that is rich in proline (Fig. S4C). Deletion of the C-terminal tail slightly slows yeast cell growth ^57^. The proline residue is often observed in mediating dynamic protein interactions ^58^. To examine whether the C-tail of Tim17 is involved in the recognition of polypeptide substrates, a yeast strain with C-tail deletion was generated (Tim17ΔC) and subjected to similar photo-crosslinking assays. The results showed that the crosslinking between Tim17 and residues 144- 159 of the substrate indeed disappeared (Fig. S4D), confirming the interactions between the C-tail of Tim17 and the middle segment of the substrate in the IMS. Thus, the N-terminal residues 128-142 of the polypeptide substrate, which remained crosslinking in the Tim17ΔC mutant, likely interact with the TM region of Tim17 and span the inner membrane (Fig. 3E).

Another set of strong bands had a molecular weight of ∼70 kDa, immediately above the substrate bands and co-appearing with the Tim17 crosslinked bands (Figs. 3A, S4B). It was likely from a TIM23 subunit with a weight of less than 10 kDa. Among the TIM23 subunits, Mgr2 has previously been shown to crosslink to the substrate and has an apparent molecular weight of 7 kDa ^49^. To examine whether these were Mgr2 crosslinked bands, an HA tag was fused to Mgr2 in the yeast cells. Indeed, crosslinking was detected between Mgr2 and the N-terminal segment of the polypeptide substrate (Fig. S4E).

Although Mgr2 plays a crucial role in stabilizing the TOM-TIM23 supercomplex *in vitro* ^49^, it is not an essential subunit like Tim17 for protein import ^48^. To examine if the crosslinking remains without Mgr2, we conducted photo-crosslinking experiments using mitochondria from Mgr2 deletion strains. While the supercomplex was not detected without Mgr2 (Fig. S4F), the polypeptide substrate could still crosslink to Tim17 (Fig. S4G). Taken together, these data confirm that the substrate spans the TOM complex and the TIM23 complex in the supercomplex (Fig. 3E). Tim17 and Mgr2 are the two major subunits of the TIM23 complex that are in close proximity to the polypeptide substrates, with Mgr2 being nonessential.

It is surprising that the Tim23 subunit is not detected in the substrate-crosslinked bands as Tim23 is considered the major translocase subunit of TIM23. In an attempt to capture substrate crosslinking to Tim23, we tagged Tim23 in the yeast cells and affinity purified the supercomplex by Tim23 pull-down. Immunoblotting of Tim23 did not reveal any evident crosslinking bands, whereas immunoblotting of the substrates displayed similar crosslinking patterns to the results above (Fig. S5A). To exclude the possibility that the interactions between Tim23 and the substrate were lost during purification, mitochondria containing accumulated supercomplexes were irradiated with UV prior to Tim23 pull-down. Crosslinking bands were still not detected by immunoblotting of either Tim23 or the substrate (Fig. S5B). By contrast, pull-down of Tim17 or Mgr2 after UV irradiation of mitochondria yielded clear crosslinking of the substrates with Tim17 or Mgr2 (Fig. S5C), similar to the crosslinking observed in the purified supercomplexes (Fig. S4B, E). These data suggest that Tim23 may not be as close to the substrate as Tim17 and Mgr2 in the supercomplex.

### Core translocase subunits in the inner membrane

The crosslinking data suggest that Tim17 and Mgr2 are likely part of the core translocase of the TIM23 complex that holds the polypeptide substrates in the supercomplex. Previous studies have shown that Tim23 is also essential for protein translocation by the TIM23 complex ^59, 60^. Since the cryo-EM density maps of the supercomplexes did not resolve individual subunits in the inner membrane, we turned to AlphaFold2 to generate models of the core translocase of the TIM23 complex ^61^, including Tim23 and Tim17, and possibly Mgr2. Modeling of Tim23 or Tim17 oligomers using AlphaFold2 did not yield stable conformations (See Methods for modeling details). Instead, a stable heterotrimer of Tim23, Tim17, and Mgr2 could be modeled with a stoichiometry of 1:1:1 (Fig. S6A). The heterotrimer model has the characteristic X shape of the core TM density of TIM23 (Fig. S6B) and could fit well into the cryo-EM map (Fig. S6C). The four TMs of both Tim17 and Tim23 form curved surfaces similar to those of the Tim22 subunit in the TIM22 complex ^62, 63^. Tim17 and Tim23 are in the “back-to-back” arrangement with the convex surfaces next to each other (Fig. 4A, B). The concave surface of Tim17 is sealed by the two TMs of Mgr2, whereas the concave surface of Tim23 is open and exposed to lipids. Thus, Tim17 and Mgr2, but not Tim23, form a channel-like structure in which lipids are excluded. The heterotrimeric model of Tim23-Tim17-Mgr2 seems to be difficult to reconcile with the idea that Tim23 is the channel-forming unit for protein translocation. However, this model is supported by large-scale deep learning-based protein-protein interaction modeling ^64^ and a cryo-EM structure of the Tim17-Tim23-Tim44 complex ^29^.

**Fig. 4.**
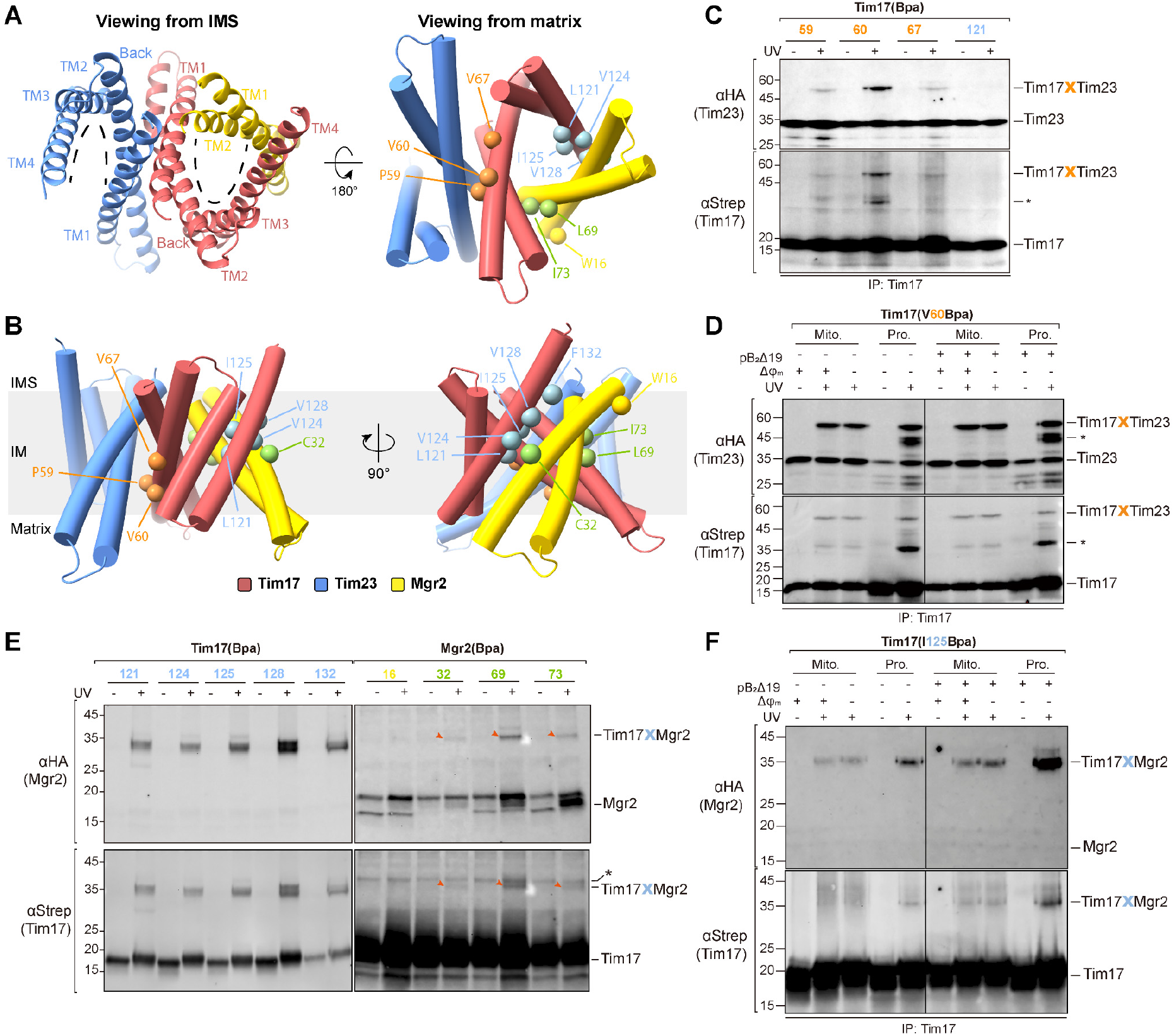
Heterotrimeric conformation of Tim23, Tim17, and Mgr2. (**A**) Top views of the Tim23- Tim17-Mgr2 heterotrimer generated by AlphaFold2. Left panel, ribbon diagram of the trimer, viewed from the IMS. Tim23, Tim17, and Mgr2 are colored Dodger blue, Indian red, and yellow, respectively. TMs are labeled. The dashed curves indicate the concave space and the openings of Tim23 and Tim17. Right panel, cylinder representation of the trimer, viewed from the matrix. The residues mutated for photo-crosslinking are shown as color-coded balls: orange, crosslinking residues of Tim17 to Tim23; light blue, crosslinking residues of Tim17 to Mgr2; light green, crosslinking residues of Mgr2 to Tim17; yellow, non-crosslinking control. The terminal flexible segments of the subunits are omitted from the diagram. (**B**) Side views of the Tim23-Tim17-Mgr2 heterotrimer, viewed from two angles differing by 90°. (**C**) Photo-crosslinking of Tim17 and Tim23. The supercomplexes were assembled with HA tagged Tim23 and Strep-His tagged Tim17 in yeast strains yLS321/pLSA1-X. Bpa was incorporated at different positions of Tim17. Tim17 was pulled down using the Ni resin after UV irradiation of mitochondria. The crosslinking products between Tim17 and Tim23 were detected by immunoblotting of the HA tag (Tim23, upper panel) and the Strep tag (Tim17, lower panel). The unidentified crosslinking bands are marked by *. (**D**) Photo-crosslinking of Tim17 and Tim23 under different conditions. Bpa was incorporated at residue 60 of Tim17. The membrane potential was dissipated by the addition of valinomycin. Pulldown and detection were performed similarly to those in **C**. Different conditions are indicated. The unidentified crosslinking bands are marked by *. (**E**) Photo-crosslinking of Tim17 and Mgr2. The supercomplexes were assembled with HA tagged Mgr2 and Strep-His tagged Tim17. Bpa was incorporated at different positions in Tim17 (left panel, yeast strains yLS411/pLSA1-X) or Mgr2 (right panel, yeast strains yLS221/pLSM1-X). Tim17 was pulled down using the Ni resin after UV irradiation of mitochondria (left panel) or pulled down first and then subjected to UV irradiation (right panel). The Ni eluents were then subjected to HA pull-down before immunoblotting. The crosslinking bands are marked by red arrows. Non-specific bands are indicated by *. (**F**) Photo-crosslinking of Tim17 and Mgr2 under different conditions. Bpa was incorporated at residue 125 of Tim17. UV irradiation was applied either on intact mitochondria (Mito.) or on digitonin-solubilized mitochondria (Pro.). Tim17 was pulled down using the Ni resin. Non-specific bands are indicated by *.

To examine whether the Tim17-Tim23-Mgr2 heterotrimer exists in mitochondria, especially during substrate import, Bpa was introduced into the TM residues of Tim17 to probe its neighboring subunits in the inner membrane (Fig. 4A, B). The mitochondria containing accumulated supercomplexes were exposed to UV and then Tim17 was pulled down. Consistent with the heterotrimer model, residues 59, 60, and 67 of TM2 at the convex surface of Tim17 could crosslink to Tim23, whereas residue 121 of TM4 at the edge of the concave surface did not show crosslinking (Fig. 4C). The “back-to-back” conformation of Tim17-Tim23 was relatively stable. As demonstrated by the V60Bpa mutant of Tim17, the crosslinking between the two subunits was maintained without accumulated substrate, membrane potential, or lipid membrane environment (Fig. 4D).

Residues 121, 124, 125, 128, and 132 at the edge of the concave surface of Tim17 were shown to crosslink to Mgr2 (Fig. 4E). Similarly, the TM residues 32, 69, and 73 of Mgr2 that are oriented toward Tim17 in the heterotrimer model were shown to crosslink to Tim17, while residue 16 of Mgr2 that is oriented away from Tim17 in the model displayed no crosslinking to Tim17 (Fig. 4E). The crosslinking efficiency between Tim17 and Mgr2 increased slightly when in the presence of the arrested polypeptide substrate (Fig. 4F), consistent with the previous results showing that Mgr2 is dynamically associated with the TIM23 complex ^49^. Interestingly, the crosslinking was greatly enhanced in the protein solution of digitonin-solubilized mitochondria.

### Translocation pathway in the inner membrane

The mapping of the polypeptide substrate in the supercomplex and the heterotrimeric Tim17- Tim23-Mgr2 model raised the possibility that the polypeptide may pass through a channel-like structure formed by Tim17 and Mgr2 in the supercomplex but not via Tim23. To explore this scenario, Bpa was introduced to the TM region of Tim17 and Mgr2 to capture the translocating polypeptide substrate (Fig. 5A, B). Two clusters of residues showed crosslinking to the substrate (Fig. 5C, D). One cluster is at the entrance of the channel on the IMS side, including residues 76, 80, and 91 of Tim17 (Fig. 5B). The other cluster is in the middle of the channel, including residues 65, 68, 121, 122, and 125 of Tim17 and residues 33 and 67 of Mgr2. Residues 121 and 125 in TM4 of Tim17 were found to crosslink to both Mgr2 and the substrate (Figs. 4E, 5C). These crosslinking results are well consistent with the Tim17-Mgr2 model since the two residues are located on the edge of the Tim17 concave surface next to Mgr2 and their side chains are oriented toward the center of the channel (Fig. S6D). These crosslinking data indicated that Tim17 and Mgr2 likely form the translocation pathway of the polypeptide substrate in the TIM23 complex. In addition to the residues in the TM region, residue 156, the second to the last residue in the C-tail of Tim17, was also introduced with Bpa to capture the substrate. The positive crosslinking band confirmed that the C-tail of Tim17 is indeed in direct contact with the substrate in the IMS (Fig. 5C).

**Fig. 5.**
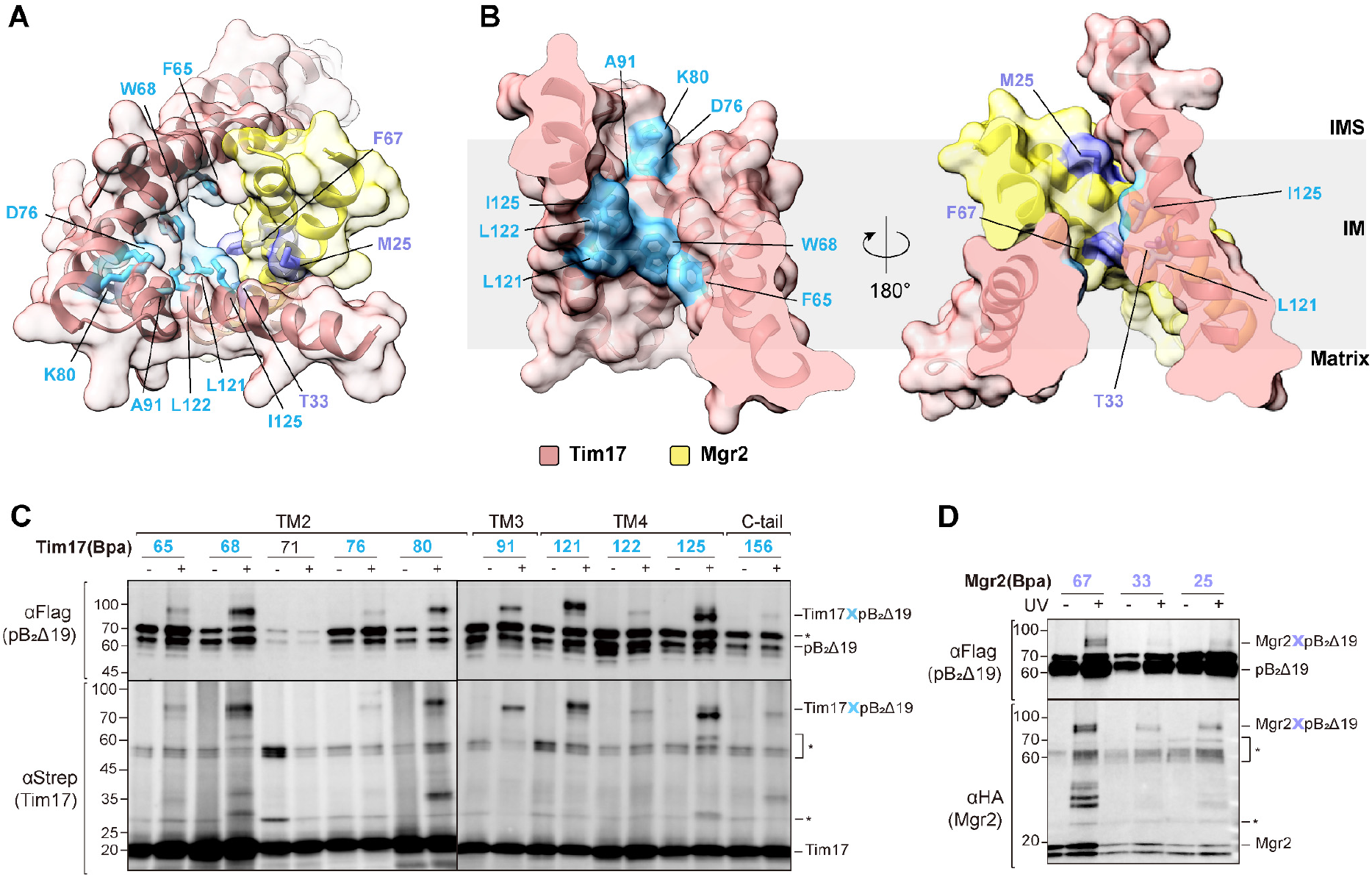
Substrate-interacting residues of the Tim17-Mgr2 heterodimer. (**A**) Top view of the Tim17-Mgr2 heterodimer. Tim17 and Mgr2 are shown in ribbon and outlined by a transparent surface. Tim17 is colored Indian red and Mgr2 is colored yellow. The substrate crosslinking residues of Tim17 and Mgr2 are labeled and shown as cyan sticks and purple sticks, respectively. (**B**) Cutaway side views of Tim17-Mgr2. The two views differ by a rotation of 180 °, showing the substrate crosslinking residues inside the import pathway. (**C**) Photo-crosslinking of Tim17 and the substrate. The supercomplexes were assembled with Strep-His tagged Tim17 in yeast strains yLS230/pLSA1-X. Bpa was incorporated at different positions in Tim17. The crosslinking products between Tim17 and the substrate were pulled down by using the Ni and Flag resin sequentially after UV irradiation of mitochondria. The residue numbers are colored according to the code in **A**. The non-crosslinking position (71) is in black. Non-specific bands are indicated by *. (**D**) Photo-crosslinking of Mgr2 and the substrate. Bpa was incorporated at different positions in Mgr2 in yeast strains yLS221/pLSM1-X. The supercomplexes were purified by pulling down Tim17 and the substrate before UV irradiation. The crosslinking products between Mgr2 and the substrate were detected by immunoblotting. Non- specific bands are indicated by *.

The mapping results of the polypeptide substrate in the supercomplex indicated that the N-terminal residues 128-142 of the substrate were captured in the import pathway in the inner membrane. Thus, the TM residues of Tim17 and Mgr2 identified above were likely photo-crosslinked to the N-terminal segment of the substrate. To test this, a cysteine residue was introduced to W68 of Tim17 (W68C) in the middle of the import pathway, as well as to the N terminal residues of the polypeptide substrate. The results showed that in the supercomplexes assembled with the cysteine mutants, Tim17(W86C) formed disulfide bonds with pB_2_-DHFR-sfGFP(135C) and pB_2_-DHFR- sfGFP(139C) (Fig. S6E), confirming the substrate and import pathway mapping results from photo-crosslinking.

The surface charge and hydrophobicity analyses of Tim17 and Mgr2 show that the two clusters of the substrate-crosslinking residues are within two distinct patches in the import pathway, a negatively charged patch of Tim17 at the entrance and a hydrophobic patch in the middle (Fig. 6A, B). Both of the patches are highly conserved. Actually, the residues of Tim17 and Mgr2 along the import pathway are all well conserved (Fig. 6C). To identify the residues that are critical for protein translocation in the pathway, highly conserved residues in the two patches were selected for mutagenesis studies, including D17, D76, and E126 from the negatively charged patch, and F65, W68, L121, L122, I125 in the hydrophobic patch. The negatively charged residues were mutated to A or K to examine how charge changes might affect protein import and cell growth. The hydrophobic residues were mutated to N, K, or D to test if hydrophobicity was important in maintaining the protein import pathway. As Mgr2 is nonessential, these mutations were also examined in the absence of Mgr2.

**Fig. 6.**
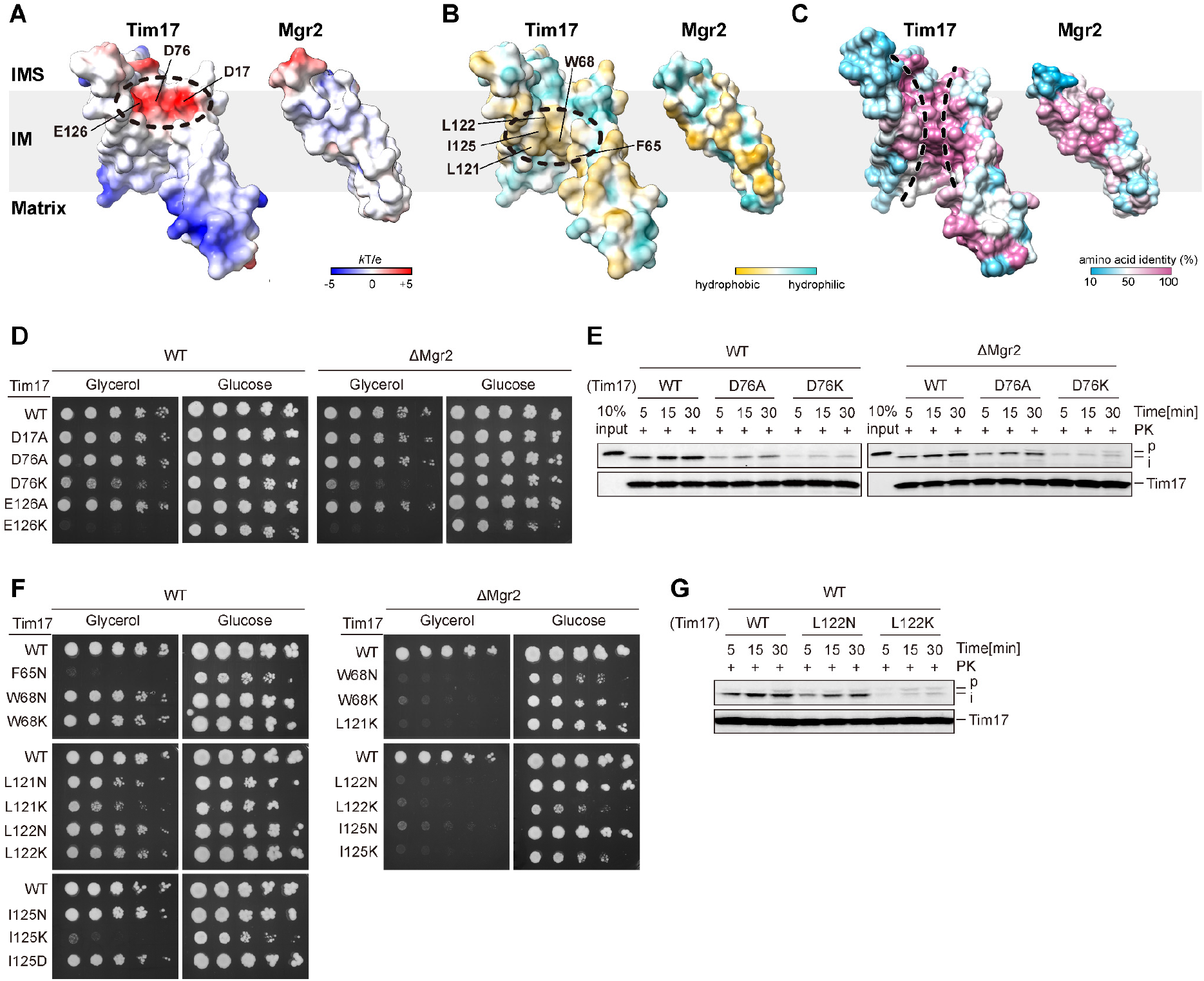
Import path of the substrate in the Tim17-Mgr2 heterodimer. (**A**) The open-book view of Tim17-Mgr2 to show surface electrostatics in the protein import pathway. The negatively charged patch is outlined by a dashed circle. The negatively charged residues are labeled. (**B**) The hydrophobicity distribution of Tim17-Mgr2, the same view as in **A**. The hydrophobic patch is outlined by a dashed circle. (**C**) Sequence conservation mapped to the Tim17-Mgr2 model. The protein import pathway is outlined by dashed lines. (**D**) Yeast spot assays to examine the growth of the mutations in the negatively charged patch. The carbon sources and the strains with and without Mgr2 are labeled. (**E**) Protein import assays, showing the import efficiency of mitochondrion mutants. A representative residue in the negatively charged patch, D76, is chosen for the assays. pB_2_167Δ19- DHFR is used as the substrate and immunoblotted by an anti-His antibody. The preprotein bands are marked by “p”. The bands of the imported substrates are marked by “i”. The amounts of mitochondria used in the reactions are indicated by Tim17 immunoblotting. (**F**) Yeast spot assays to examine the growth of the mutations in the hydrophobic patch. (**G**) Protein import assays. A representative residue in the hydrophobic patch, L122, is chosen for the assays.

In the negatively charged patch, the yeast growth assays showed that the A mutations had little effect on cell growth, whereas the K mutations caused substantial growth defects (Figs. 6D, S6F). The D17K mutant could not be obtained from 5-fluoroorotic acid selection (Fig. S6F), probably due to severe mitochondrial dysfunction caused by the mutation. The E126K mutant displayed slow growth on the fermentable carbon source (glucose), and no growth on the non-fermentable carbon source (glycerol) (Fig. 6D). Only the D76K mutant could grow on both glucose and glycerol, but with obvious growth defects (Fig. 6D). Furthermore, the mitochondrial import assays revealed that D76A reduced protein import efficiency to ∼ 40%, while D76K significantly impaired protein import, with only ∼20% efficiency of the wild type (Fig.6E). Therefore, the highly negatively charged patch comprising of three D/E residues plays a critical role in protein translocation through the inner membrane. An alanine mutation that moderately reduces charges is relatively tolerable, but the introduction of positive charges from lysine could be detrimental to protein translocation. Notably, these mutants had similar growth phenotypes and mitochondrial import efficiencies in the absence of Mgr2 (Fig. 6D, E). This is consistent with the Tim17-Mgr2 model in which Mgr2 does not have obvious contribution to the charge of the import pathway (Fig. 6A).

In the hydrophobic patch, mutations to the polar/charged residues resulted in varying degrees of impairment to cell growth. Residue F65 was particularly sensitive to mutation. F65K and F65D could not be obtained from selection (Fig. S6F), while F65N showed no growth on glycerol (Fig. 6F). Most D mutants in the hydrophobic patch were intolerable, except for I125D (Figs. 6F, S6F).

Generally, K mutations had a greater effect on cell growth than N mutations in most sites, such as L121, L122, and I125 (Fig. 6F). Consistent with the growth defects, L122K impaired protein import to a much greater extent than L122N (Fig. 6G). Striking growth defects were observed when the mutations were combined with Mgr2 deletion, with essentially no mutants able to grow on glycerol (Fig. 6F). Growth on glucose was also significantly impaired. Apparently, the effect of Mgr2 deletion on the polar/charge mutations of the hydrophobic patch differed from its effect on the mutations of the negatively charged patch. These results support the Tim17-Mgr2 model, in which Mgr2 shields the hydrophobic patch of Tim17 from lipids in the channel-like structure (Fig. 6B), thus the polar/charge mutations of the hydrophobic residues could be relatively tolerated. When Mgr2 is absent, the hydrophobic patch of Tim17 in the middle of the import pathway is exposed to the lipid environment. Introduction of polar/charge residues in the patch would be much less tolerated in lipids, resulting in an impaired protein import pathway and mitochondrial dysfunction.

The photo-crosslinking and mutagenesis data together revealed a protein import pathway in the inner membrane that is formed by Tim17 and Mgr2. The efficient translocation of proteins through this pathway is dependent on both a negatively charged patch at the entrance and a hydrophobic patch located in the middle of the pathway. Tim17 plays a vital role in providing the essential structural elements required for the pathway, while the Mgr2 subunit, which is nonessential, contributes to the formation of a channel-like structure that shields and monitors the polypeptide substrates during translocation.

## Discussion

### Assembly of the TOM-TIM23 supercomplex

The use of the arrested polypeptide substrates has been crucial in biochemical characterization of the TOM and TIM23 complexes, particularly when combined with import assays *in vitro*. To facilitate structural characterization, we designed a series of polypeptide substrates for the *in vivo* assembly of the TOM-TIM23 supercomplex. The cryo-EM structures of the supercomplexes revealed that one of the two Tom40 channels in the TOM complex is engaged in protein import.

However, this may be due to steric hindrance at the cytosolic side caused by the bulky sfGFP in the designed substrates. It is unclear if both Tom40 channels may actively translocate natural preprotein substrates simultaneously. In addition, it remains uncertain if the angle and distance between the TOM and TIM23 complexes, as shown in the structures, accurately reflect the relationship between the two complexes in mitochondria during protein import. While the distance of ∼5 nm is consistent with a previous result from chemical crosslinking/mass spectrometry ^65^, a cryo-electron tomography study at a low resolution suggested a larger distance between the outer and inner membranes at the import sites ^66^.

### The TOM complex in the supercomplex

The TOM complex does not appear to undergo significant conformational changes during protein import. In particular, the Tom40 pore shows minimal changes, with an r.m.s.d. of ∼ 1.0 Å between the structures of the substrate-engaged Tom40 pore and the idle Tom40 pore ^16^. Indeed, the Tom40 pore has a minimal diameter of 11Å, which is more than enough to accommodate an extended polypeptide during translocation.

The structure of the substrate-engaged TOM complex reveals a “glutamine-rich” patch (patch 2) that mediates substrate interaction in the Tom40 pore. This patch is relatively conserved among different species, with Q99 and Q156 being highly conserved and Q128 being not (Fig. S7A). R140 in yeast Tom40 aligns with N and Q residues in metazoan species (Fig. S7A). M94 is located in an unconserved loop of yeast Tom40, however, in the structure, it is close to the conserved N111 residue of human Tom40 (Fig. S7B). These observations suggest that the “polar-rich” patch is a common feature across species for facilitating substrate translocation. In contrast to the “polar-rich” patch revealed by the extended and channel-spanning polypeptide substrate in this study, an earlier investigation revealed two other substrate interacting patches of Tom40 using a positively charged pre-sequence (interacting with patch 1) and a hydrophobic carrier protein (interacting with patch 3) that were stuck inside the Tom40 pore ^13^. Consistently, patch 1 is positioned close to negatively charged residues in Tom40 which could guide pre-sequences through the pore ^13^. Patch 3 comprises hydrophobic residues that may help to stabilize hydrophobic substrates during import.

The folded sfGFP molecule is stuck on the cytosolic side of Tom40 as an import blocker in the supercomplex structure. While not in a fixed orientation in the structure, the density map shows that sfGFP is likely to have direct contact with the cytosolic loop between β14 and β15 (L14-15) of Tom40 (Fig. S7C). The loop is disordered in the idle TOM complexes ^14, 16^. The tip of the loop is rich in hydrophobic residues which may help to stabilize and align the unfolded polypeptide substrates prior to their entrance into the central pore (Fig. S7D).

### The TIM23 complex in the supercomplex

The TIM23 complex is highly dynamic ^33, 34^. The exact organization of the TIM23 complex was unclear, especially during substrate import. The IMS domains of Tim50, Tim21, and Tim23 have been shown to interact with each other and with substrates ^30, 31, 67, 68^. On the matrix side, Tim44 interacts with Tim23, Tim17, and substrates ^36, 37^. The purified supercomplexes for cryo-EM studies essentially contain all TIM23 subunits as judged by mass spectrometry. However, the cryo-EM density maps show little density on the IMS and matrix sides, suggesting that the soluble domains of these subunits are flexible and/or the interactions between subunits are highly dynamic. Consistently, previous biochemical studies suggest that Tim44, Tim50, and Hsp70 are not stably associated with the supercomplex ^52, 53^.

The strong density of the TIM23 complex is only observed in the trans-membrane region. Previous chemical crosslinking provided limited details regarding the interactions between the transmembrane regions of the TIM23 complex, likely due to the lack of reactive residues in the TMs, such as Lys. Guided by Alphafold2 modeling, we demonstrated the presence of the Tim17-Tim23- Mgr2 heterotrimer in mitochondria by site-specific photo-crosslinking. Nevertheless, this heterotrimer organization could be dynamic during protein import. For example, Mgr2 is nonessential for substrate import, but is critical for supercomplex formation ^49^. Thus, Mgr2 may not be stably associated with Tim17 in the idle state, but could form a dynamic channel-like structure with Tim17 to aid substrate import in the active state. Similarly, Tim23 has been shown to form voltage-gated channels ^26^. TM2 of Tim23 has an aqueous-facing helical surface, the exposure of which is affected by membrane potential and substrate import ^69, 70^. These properties suggest that Tim23 may have other conformations that are not present in the heterotrimer.

### Polypeptide import pathway in the TIM23 complex

The ambiguity of the core translocase unit in the TIM23 complex has been largely due to the lack of direct evidence showing the immediate surrounding environment of the hydrophilic polypeptide substrates passing through the hydrophobic lipid membrane. By stably trapping the substrate pB_2_167Δ19-DHFR-sfGFP in the supercomplex, we were able to employ photo-crosslinking to precisely map the polypeptide substrate in the inner membrane and examine the subunits along the import pathway. Unexpectedly, the polypeptide segment is surrounded by Tim17 and Mgr2 in the supercomplex, but not by the generally expected Tim23 subunit. In line with the finding, some species have been shown to have only the Tim17 subunit, but not Tim23, in mitochondria ^71–73^. The Tim17- directed protein import is also implicated by a study on stendomycin, a compound that inhibits mitochondrial protein import by directly targeting the TIM23 complex ^74^. Single point mutations on Tim17 were found to be sufficient to confer resistance to stendomycin, underscoring the critical role of Tim17 in controlling protein import.

As preprotein substrates pass the inner membrane via Tim17, Mgr2 seals the lateral opening of Tim17 to the lipid membrane by forming a channel-like structure, consistent with its function as a lateral gatekeeper for preprotein sorting. In this conformation, Tim17 and Mgr2 could tightly hold the polypeptide substrates, explaining the observation that Mgr2 is required for the supercomplex formation ^49^. In the middle of the import pathway, from the hydrophobic patch, four highly conserved residues F65, W68, and L121 of Tim17 and F67 of Mgr2 form the narrowest restriction of the pathway (Fig. S8A-D). The restriction has a diameter of ∼7-8 Å (Fig. S8A), similar to the opening of the hydrophobic pore ring in the SecY channel during protein translocation. The opening size could allow passage of an extended polypeptide and prevent leakage of small ions ^75^. After passing the restriction region, the channel-like structure becomes wide open on the matrix side. The TMs of Tim17 and Mgr2 on this side actually extend outwards and do not maintain an enclosed channel (Fig. S8B). This conformation may facilitate the delivery of the substrate to the receptor Tim44 on the matrix side of the TIM23 complex.

While Mgr2 makes direct contact with polypeptide substrates in the TIM23 complex, it is not an essential subunit for protein import. Without Mgr2, the TOM-TIM23 complex cannot be stabilized *in vitro* possibly because Mgr2 is not present to hold the polypeptide substrates to Tim17 once in solution. However, our experiment showed that polypeptide substrates can still crosslink to Tim17 while in the inner membrane. This suggests that Tim17 alone, without channel formation, may be sufficient to facilitate polypeptide translocation. In this scenario, the protein import pathway primarily consists of the concave surface of Tim17 that is exposed to the lipids. The negatively charged patch at the entrance of the import pathway may distort the local lipid bilayer structure, causing membrane thinning to facilitate protein translocation. Similar mechanisms have been proposed for other protein translocation systems, including the YidC insertase ^76^, the ER membrane complex ^77, 78^, the TIM22 complex ^63^, and ER retro-translocation mediated by Hrd1 ^79^. However, it remains possible that, in addition to a single copy of Tim17, more copies of Tim17 or other subunits, such as Tim23, may be involved in the formation of the protein conducting channel at different stages of protein translocation. In either case, Tim17 is likely to be the key translocase subunit along the import pathway to facilitate the passage of preproteins through the inner membrane.

In summary, we established a methodology to assemble and stabilize the TOM-TIM23 supercomplex for comphensive structural and biochemical analyses. The cryo-EM structures of the supercomplex show that the TOM and TIM23 complexes could be physically connected for efficient import of protein substrates through the mitochondrial double membranes. While passing through the outer membrane, the polypeptide substrates run along a glutamine-rich path in the inner wall of the Tom40 channel (Fig. S9). In the inner membrane, the polypeptides are held by a channel-like structure formed by Tim17 and Mgr2 (Fig. S9). Taken together, we mapped the pre-sequence import pathway in detail, revealed an unexpected Tim17-Tim23-Mgr2 heterotrimer, and highlighted the critical role of Tim17 as a translocase subunit directly mediating substrate translocation through the inner membrane.

## Materials and Methods

### Yeast strains and plasmid construction

For structural studies, the *S. cerevisiae* strain yLS110 was generated from the W303 strain by fusing a His_10_ tag to the C-terminus of Tom22 in the genome. For substrate expression, the first 167 amino acids of cytochrome b2 with residues 47-65 deleted were fused to a C-terminal sfGFP and a twin-StrepII tag (pB_2_167Δ19-sfGFP-twinStrepII). DNA encoding pB_2_167Δ19-sfGFP-twinStrepII was inserted into the pRS423 vector downstream of the inducible galactose promoter GAL1, resulting plasmid pLSB1. The yLS110 strain was transformed with pLSB1 and selected on yeast synthetic defined medium without Ura and His (SD-Ura/His) to yield the supercomplex expressing strain yLS111. Similarly, the strain expressing the supercomplex assembly with pB_2_167Δ19-DHFR-sfGFP was generated using the yLS110 strain and the pLSB2 plasmid in which DHFR was inserted between residues 125 and 126 of pB_2_167Δ19-sfGFP-twinStrepII.

For photo-crosslinking assays, Tom40, Tim17, Tim23, and Mgr2 were either N- or C-terminally tagged in the genome or expressed with the tags from the pRS31X series of vectors in the knock-out strains. The photoreactive unnatural amino acid *p*-benzoyl-L-phenylalanine (Bpa) was introduced by transformation of a plasmid encoding a modified tRNA-synthetase that charges tRNA with Bpa and the tRNA suppresses the amber stop codon ^80^.

### Cryo-EM data acquisition

The purified supercomplex was concentrated to 5-7 mg/mL. For the supercomplex assembled with pB_2_Δ19-DHFR-sfGFP, 5 μM methotrexate was added and incubated on ice for 30 min before cryo sample preparation. A 2.5-3 μL aliquot of protein sample was dropped onto the glow discharged holey carbon grids (Quantifoil Au R1.2/1.3, 300 mesh), blotted with a Vitrobot Mark IV (ThemoFisher Scientific) (1.5 s blot time, 5 s wait time, 100% humidity at 6 °C), and plunge-frozen in liquid ethane cooled by liquid nitrogen.

The cryo-EM grids were imaged on a Titan Krios transmission electron microscope (FEI) at 300 kV with a magnification of 81,000 ×. Images were recorded by a Gatan K3 Summit direct electron detector using SerialEM ^81^. Defocus values varied from -1.0 to -1.2 μm. Each image was dose fractionated to 40 frames with a total electron dose of 60 e^-^Å^-2^ and a total exposure time of 3.2 s. All movie stacks were motion-corrected using MotionCor2 ^82^ and the defocus values were estimated using Gctf ^83^.

### Structure determination

To determine the structure of the supercomplex with pB_2_167Δ19-sfGFP, a total of 8,041 movie stacks were recorded. All data processing was performed using Relion ^84^. A total of 1,869 particles were manually picked and subjected to 2D classification to generate templates for automatic particle picking. A total of 2,284,465 particles were auto-picked from 6,818 images that were manually selected. The picked particles were subjected to three rounds of 2D classification, and 1,901,689 particles were selected for subsequent 3D classification. The initial model was generated from the selected particles with high quality. One class with 113,571 particles was selected from the last round of 3D classification for a new round of 2D classification and for generating a template for another round of particle auto-picking. The auto-picked 3,215,742 particles were subjected to two rounds of 2D classification. The 2D classes with features in the the TIM23 disc were selected to generate an initial model (3D-Ref1) that had transmembrane density in the TIM23 disc. Two rounds of unsupervised 3D classifications with 3D-Ref1 as the reference map were performed, followed by another round of supervised 3D classification with four reference maps, including 3D-Ref1 and three newly generated maps referred as 3D-Refset2. Two classes were merged for final refinement. The workflow of data processing is illustrated in Fig. S1C-G. The model of the TOM complex (PDB ID: 6UCU) was fitted in the density map of the large disc using Coot ^85^. Tom22 was extended to the full length in the density map on the basis of the model predicted by AlphaFold2 (https://alphafold.ebi.ac.uk/entry/P49334).

The structure of the supercomplex with pB_2_167Δ19-DHFR-sfGFP was determined using a similar process as described above (Fig. S2A-D). To improve the local resolution of the TOM complex, a mask including the TOM40 dimer and sfGFP was applied to the particles for 3D classification.

Subsequently, 3D classification without alignment was performed with a mask covering only the substrate-engaged Tom40 channel and sfGFP. A subset of 103,262 particles was further subjected to iterative cycles of CTF refinement, Bayesian polishing, and local masked 3D refinement, yielding a map of 4.10 Å resolution (Fig. S2E-H). The model of the TOM complex (PDB ID:6UCU) was fitted in the density map of the substrate engaged TOM complex using Coot ^85^. After manual building and modification, the model was refined in real space using PHENIX ^86^. The model of sfGFP (PDB ID:2B3P) was fitted in the density map and subjected to rigid body refinement using PHENIX.

### Modeling of Tim23-Tim17-Mgr2

Structure prediction was performed for yeast Tim23 (Uniprot ID: P32897), Tim17 (Uniprot ID: P39515), and Mgr2 (Uniprot ID: Q02889), using Alphafold2 on the Google Colaboratory server ^87, 88^. Various homo- and hetero-oligomers were tested, including homodimers (Tim23)_2_ and (Tim17)_2_; heterodimers Tim23-Tim17, Tim23-Mgr2, and Tim17-Mgr2; homotrimers (Tim23)_3_ and (Tim17)_3_; hetero-trimers (Tim23)_2_-Tim17, (Tim17)_2_-Tim23, and Tim23-Tim17-Mgr2; hetero-tetramer Tim23- Tim17-(Mgr2)_2_. For each test, five models were generated and evaluated based on their predicted Local Distance Difference Test (pLDDT) scores (per-residue confidence), Predicted Aligned Error (PAE) plots (domain position confidence), and consistency of the subunit orientations between parallelly predicted models (Fig. S10). Results showed that the Tim23-Tim17 heterodimer, the Tim17-Mgr2 heterodimer, and the Tim23-Tim17-Mgr2 heterotrimer consistently had high pLDDT scores in the TM regions and low PAE within and between subunits (Fig. S10C, E, J). The average r.m.s.d. between the core transmembrane regions in the models of Tim23-Tim17, Tim17-Mgr2, and Tim23-Tim17-Mgr2 all had values of ∼ 1.1 Å, suggesting each prediction yielded essentially the same conformation of transmembrane regions in all five models. The Tim23-Tim17-Mgr2 models could actually be generated by superimposing Tim17 from the Tim23-Tim17 and Tim17-Mgr2 models, demonstrating the robustness of AlphaFold2 prediction. Furthermore, human Tim23-Tim17b1-Romo1 (equivalent of yeast Mgr2) was modeled by AlphaFold2 to examine if the organization of Tim23- Tim17-Mgr2 is conserved. The resultant models had high pLDDT scores, low PAE, and the same arrangement of the three subunits as their yeast countparts in the Tim23-Tim17-Mgr2 heterotrimer (Fig. S10L).

### Photo-crosslinking assays

For photo-crosslinking in protein solutions, the supercomplex with incorporated Bpa was affinity purified by sequential pulldown of the His-tag and Flag-tag. 5 μM methotrexate was added to stabilize DHFR folding in the substrate. Half of the purified sample was irradiated by UV for 12 min on ice, and the other half was kept on ice in the dark. Samples were analyzed by SDS-PAGE and immunoblotting.

For photo-crosslinking in mitochondria, the isolated mitochondria were incubated with 20 μM methotrexate at 4°C for 30 min. Half of the sample was UV-irradiated for 30 min on ice, and the other half was kept on ice in the dark. Then, mitochondria were solubilized with 1% SDS buffer (20 mM Tris-HCl pH8.0, 150 mM NaCl, 1% SDS, 8M urea) at 98°C for 5 min. The insoluble material was removed by ultracentrifugation at 200,000 *g* for 30 min at 25°C. The supernatant was diluted 5-fold in 0.5% Triton-X100 buffer (20 mM Tris-HCl pH8.0, 150 mM NaCl, 8 M urea, 0.5% Triton-X100, 20 mM imidazole) and incubated with Ni-NTA resin for 90 min at room temperature. The bound proteins were eluted with 250 mM imidazole and analyzed by SDS-PAGE and immunoblotting.

### Disulfide crosslinking

Isolated mitochondria were resuspended in MIB buffer (10 mM Tris-HCl pH 7.4, 250 mM sucrose) and incubated with 20 μM methotrexate to stabilize DHFR folding. Then, copper phenanthroline was added at a final concentration of 1 mM to promote formation of the disulfide bond between the cysteine mutants of Tim17 (68C) and the substrate (135C, 139C). The treated mitochondria were collected by centrifugation, solubilized with 1% SDS at 98°C for 5 min. After removing the insoluble material, the supernatant was diluted 5-fold in 0.5% Triton-X100, and then subjected to Ni-NTA and Flag affinity purification. The eluted proteins were analyzed by SDS-PAGE and immunoblotting.

### Yeast spot assays with the Tim17 mutants

The pLSA1 plasmid was transformed into W303-1a or yLS430 strain. The endogenous Tim17 gene was replaced by the selectable cassette fragments to generate the Tim17 shuffling strains (yLS270 and yLS280). The Tim17 wild-type (pLSA4) and mutation plasmids (pLSA4-X) were transformed into these Tim17 shuffling strains, with the empty pRS315 plasmid as the control. The strains were selected on the synthetic medium with 5-fluoroorotic acid to eliminate the pLSA1 plasmid. To examine yeast survival and growth, cells were cultured in medium containing 3% (w/v) glycerol or 2% (w/v) glucose at 30℃ with shaking at 180 rpm. The cell culture was then diluted to an OD600 of 0.4 and subjected to a 5-fold serial dilution. The cells were spotted on the YPD or YPG medium plates and cultured at 30°C for 72h to monitor growth.

### Protein import assays

The mitochondria were incubated with precursor proteins in import buffer (250 mM sucrose, 10 mM MOPS-KOH (pH 7.2), 80 mM KCl, 5 mM dithiothreitol, 5 mM MgCl_2_, 2 mM ATP, 2 mM NADH, 3% (w/v) bovine serum albumin, 5 mM creatine phosphate and 0.1 mg/mL creatine kinase) at 25 ℃ for 5, 15 or 30 min. The import reaction was stopped by adding AVO-mix (0.8 mM antimycin A, 0.1 mM valinomycin and 2 mM oligomycin) and then incubated with 0.5 mg/mL proteinase K on ice for 30 min to remove the nonimported precursor proteins. After proteinase K treatment, 1 mM phenylmethyl sulfonyl fluoride (PMSF) was added. The samples were then washed with SEM buffer and analyzed by SDS-PAGE and immunoblotting.

## Acknowledgments

We thank Dr. Ning Gao, Dr. Xinzheng Zhang, and Dr. Ningning Li for helping with cryo-EM calculation, Dr. Qing Li for helping with yeast genetics, and the National Center for Protein Sciences at Peking University for assistance with protein purification. The cryo-EM data were collected at the cryo-EM platform of Peking University. The computation was supported by the High-Performance Computing Platform of Peking University. This work is supported by National Natural Science Foundation of China (NSFC) (31870835 to LL).

## Author contributions

X.Z. and Y.Y. prepared the protein samples for structural studies. S.W., X.Z., and Y.Y. performed the crosslinking assays. G.W., X.Z., and Y.Y. collected the cryo-EM data and determined the structures.

D.S. explored the sample preparation strategies at the beginning of the project. X.O. helped with preparation of the proteins and cryo-grids. Y.L. helped with yeast genetics. L.L. supervised the project. L.L., X.Z., Y.Y., G.W., and S.W. prepared the manuscript.

---

**Fig. S1.**
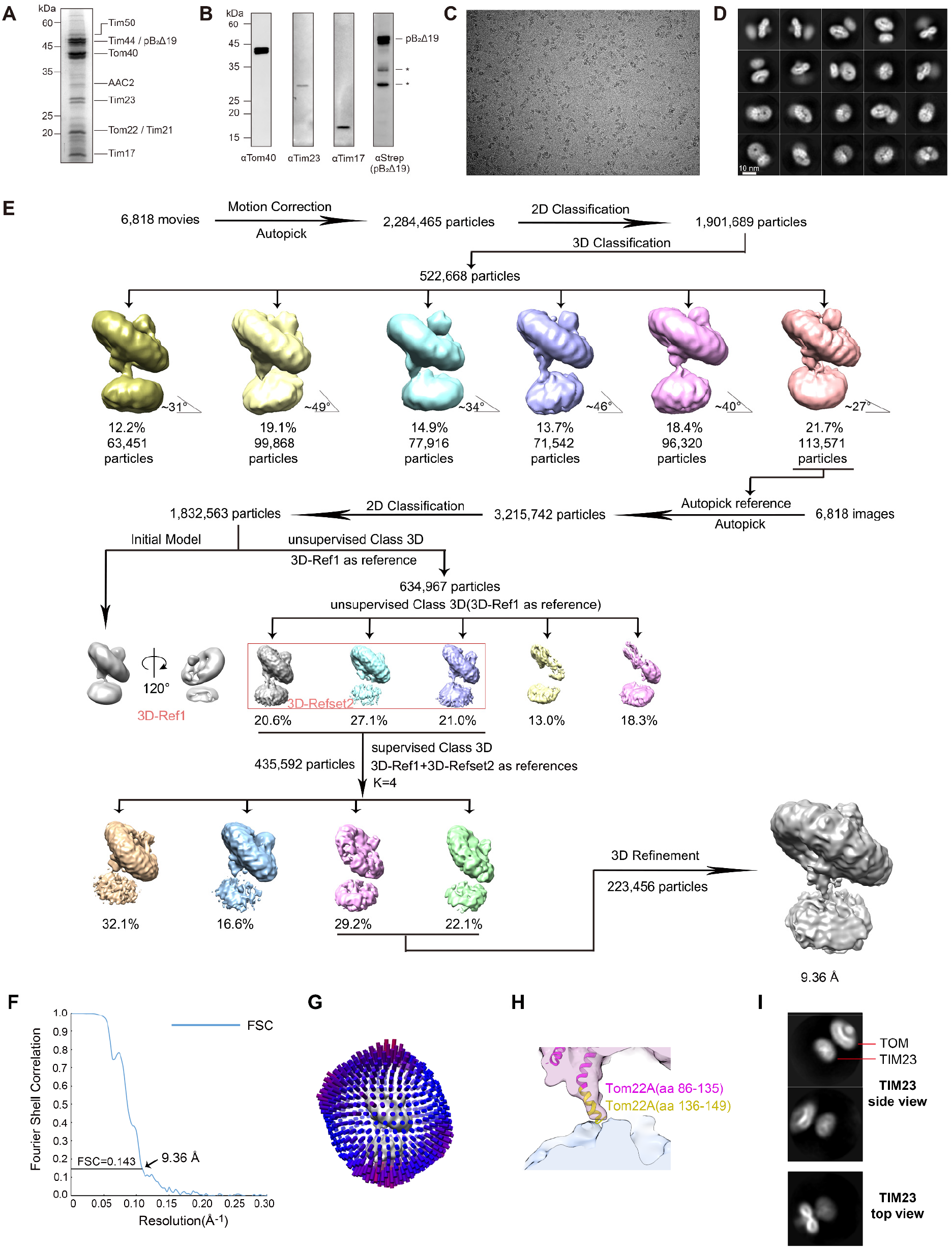
Purification and structure determination of the TOM-TIM23 supercomplex with pB_2_167Δ19-sfGFP. (**A**) Coomassie blue staining of the purified supercomplex in SDS-PAGE. The protein bands marked in SDS-PAGE have been confirmed by mass spectrometry. (**B**) Immunoblots of the substrate and the major translocase subunits in the purified supercomplex. The degradation bands of the polypeptide substrate are marked by *. (**C**) Representative raw image of the supercomplex collected by a Titan Krios with a K3 detector. (**D**) Representative 2D classes of the supercomplex. (**E**) Flow chart of data processing. The angles between the two discs in different 3D classes are labeled. See the details of data processing in Methods. (**F**) Gold standard Fourier shell correlation (FSC) curve with the estimated resolution of the final map at 0.143. (**G**) Eulerian angle distribution of the particles used for supercomplex reconstruction. (**H**) A close-up view of the connection between the TOM and TIM23 discs. The structure of the TOM complex (6UCU) is fitted in the density. The Tom22 segment in the structure (residues 86-135) is colored magenta. The model of the Tom22 C-terminal segment (residues 136-149) is from the AlphaFold Protein Structure Database and colored yellow. (**I**) Representative 2D classes of the supercomplex particles. The TIM23 disc is placed in the center of the box for better alignment. The characteristic X shape of the TIM23 TM region can be observed in the side view.

**Fig. S2.**
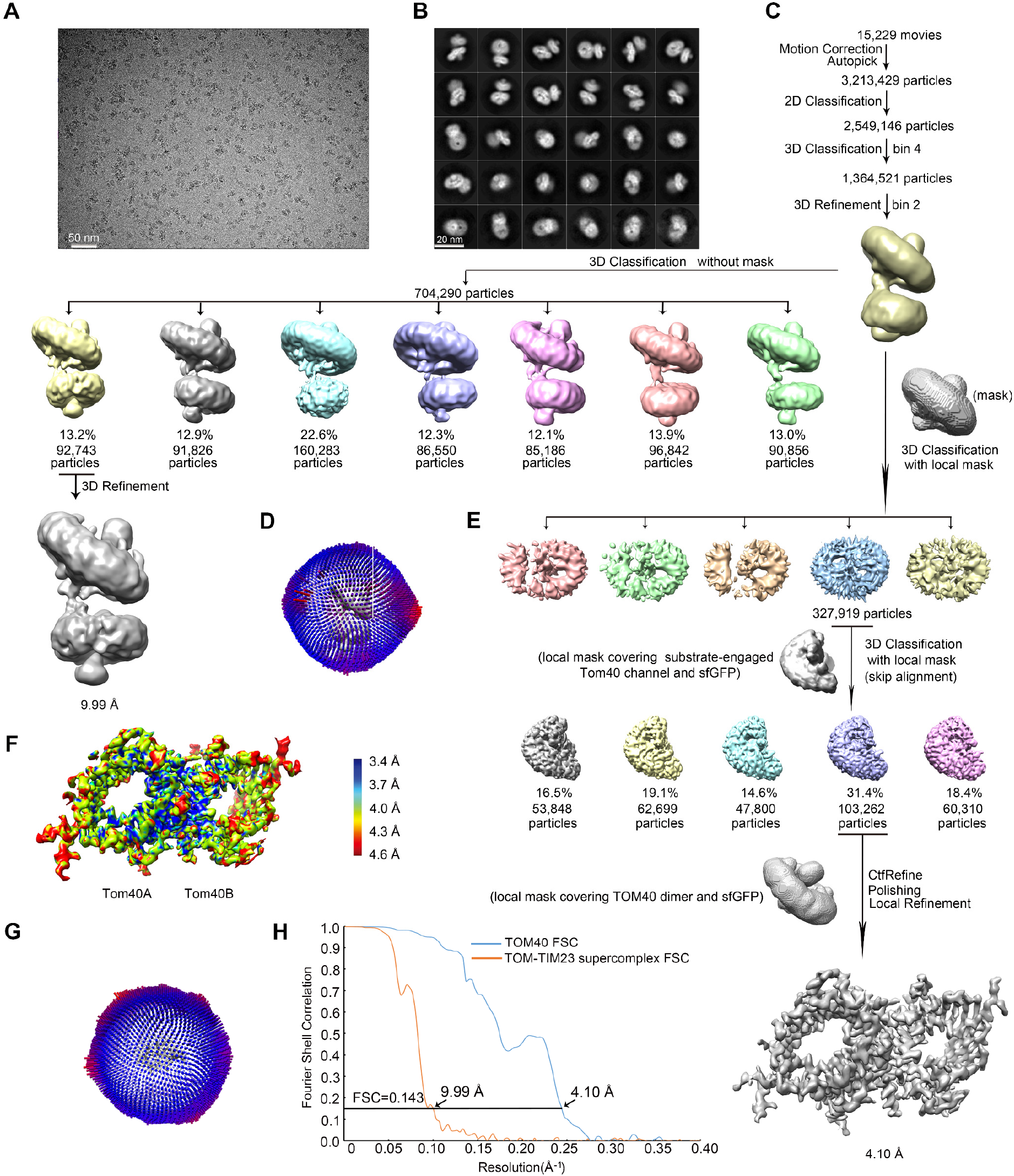
Cryo-EM 3D reconstructions of the supercomplex with pB_2_167Δ19-DHFR-sfGFP. (**A**) Representative raw image of the super-complex collected by a Titan Krios with a K3 detector. (**B**) Representative 2D classes. (**C**) Flow chart of data processing for the super-complex. (**D**) Eulerian angle distribution of the particles used for supercomplex reconstruction. (**E**) Flow chart of data processing focusing on the TOM complex. (**F**) Local resolution estimation of the TOM complex. (**G**) Eulerian angle distribution of the particles used in the local TOM reconstruction. (**H**) FSC curves with the estimated resolution at 0.143 of the supercomplex and the TOM maps.

**Fig. S3.**
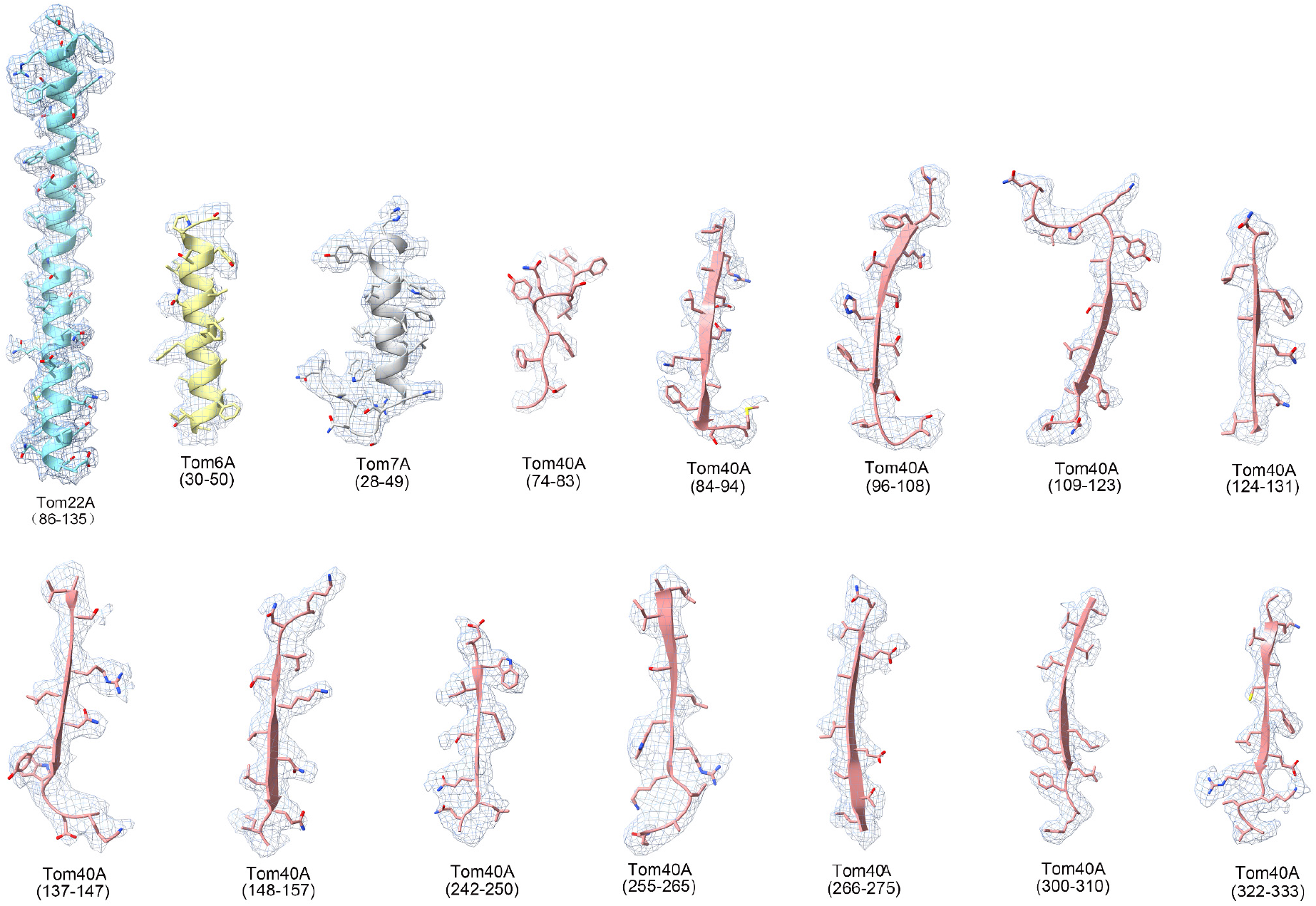
Examples of the fit of the models into density maps. Density maps (grey mesh) and the models of the selected peptide segments are shown. Residues at the beginning and end of each segment are indicated. The residue side chains are shown as sticks.

**Fig. S4.**
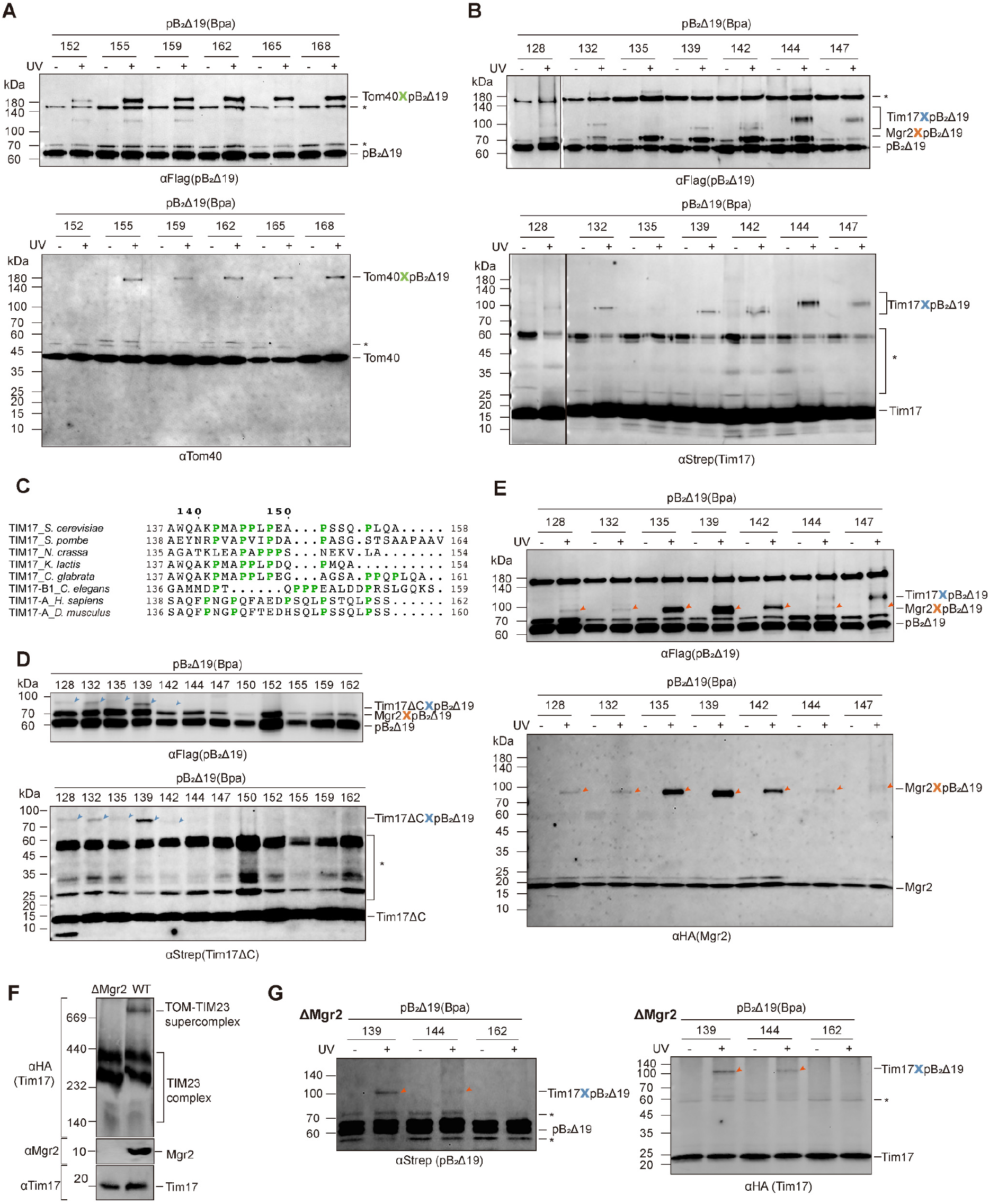
Photo-crosslinking of the polypeptide substrates to the subunits in the supercomplex. Similar crosslinking assays as described in Fig. 3, showing the controls without UV exposure. (**A**) Crosslinking between the C-terminal segment of the substrate and Tom40. The same protein samples were immunoblotted with the anti-Flag antibody to detect the substrates (upper panel) and with the anti-Tom40 antibody (lower panel). The non-specific bands in the immunoblots are marked by *. (**B**) Crosslinking between the N-terminal segment of the substrate and Tim17. The same protein samples were immunoblotted with the anti-Flag antibody to detect the substrates (upper panel) and with the anti-Strep antibody to detect Tim17 (lower panel). (**C**) Sequence alignment of Tim17 C-tail. The proline residue is highlighted in green. The multiple-sequence alignment was performed with ClustalOmega. The sequences are from *Saccharomyces cerevisiae* (*S. cerevisiae,* Uniprot P39515), *Schizosaccharomyces pombe* (*S. pombe,* Uniprot P87130), *Neurospora crassa* (*N. crassa,* Uniprot P59670), *Homo sapiens* (*H. sapiens*, Uniprot Q99595), *Mus musculus* (*M. musculus*, Uniprot Q545U2), *Caenorhabditis elegans* (*C. elegans,* Uniprot O44477). (**D**) Crosslinking between the substrate and the Tim17 C-tail deletion mutant (Tim17ΔC) from strains yLS290/pLSB4-X. Protein extraction and crosslinking were performed same as in **B**. The non-specific bands in the immunoblots are marked by *. (**E**) Crosslinking between the N-terminal segment of the substrate and Mgr2 in yeast strains yLS240/pLSB4-X. The same protein samples were immunoblotted with the anti-Flag antibody to detect the substrates (upper panel) and with the anti-HA antibody to detect Mgr2 (lower panel). The crosslinking bands are marked by red arrows. (**F**) Dependence of supercomplex formation on Mgr2. The supercomplex band is absent in BN-PAGE (upper panel) when Mgr2 is deleted. The two lower strips are from SDS-PAGE to detect Mgr2 and Tim17. (**G**) Crosslinking between the polypeptide substrate and Tim17 from the Mgr2 knockout yeast strains. The protein samples were immunoblotted with the anti-Strep antibody to detect the substrates (left panel) and with the anti-HA antibody to detect Tim17 (right panel). The crosslinking bands at the residue positions 139 and 144 are marked by red arrows. Residue 162 of the substrate is located in Tom40 and shows no crosslinking to Tim17.

**Fig. S5.**
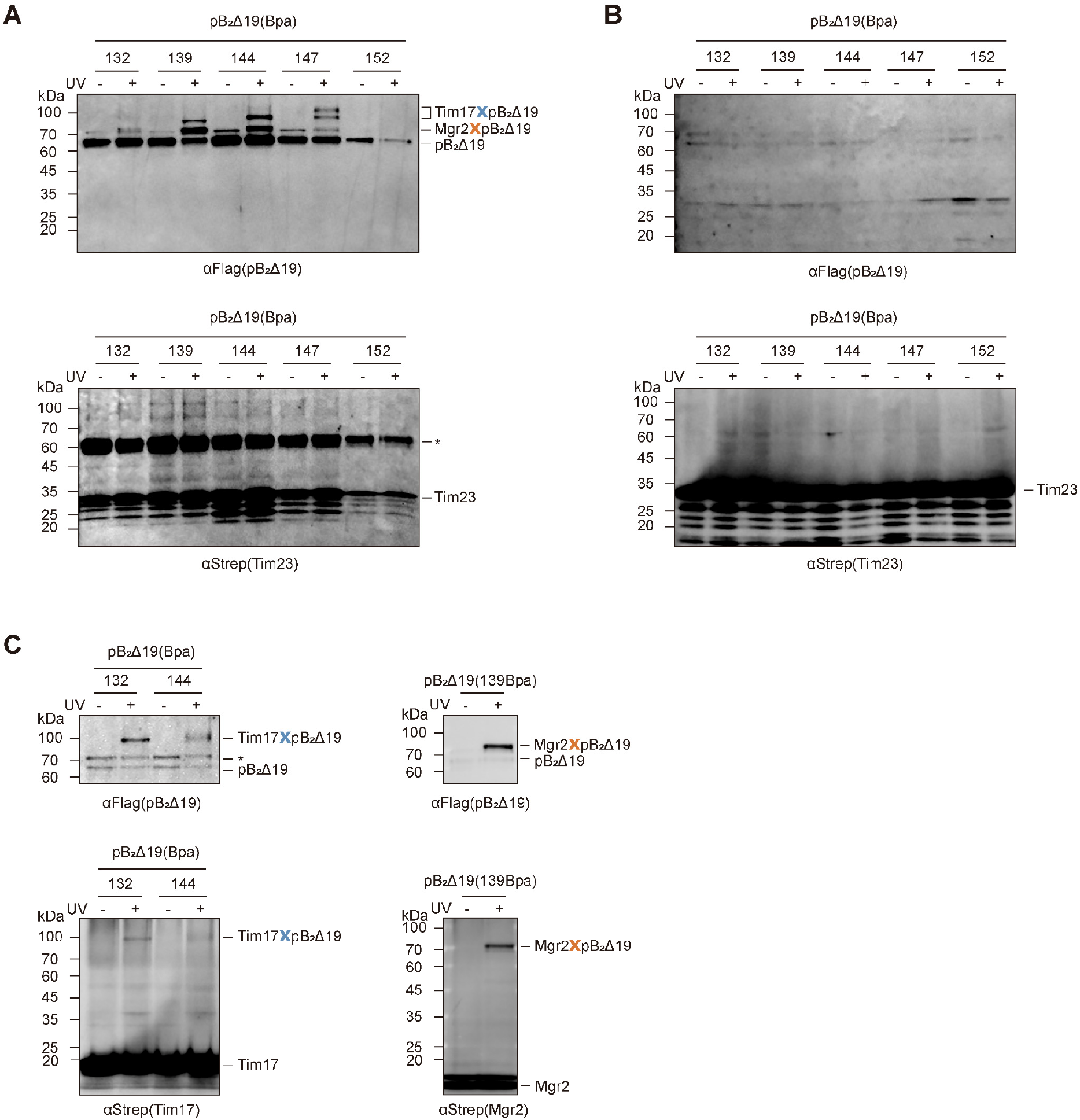
Detection of photo-crosslinking between the substrate and Tim23. (**A**) The Bpa was incorporated into the polypeptide substrate at the indicated residues in the N-terminal segment. The supercomplexes were assembled with these Bpa-incorporated substrates in yeast strains yLS310/pLSB4-X, and then purified by pulling down Tim23 (Strep-His tagged) and the substrate (Flag tagged) sequentially. The purified supercomplexes were subjected to UV irradiation, and immunoblotted with the anti-Flag antibody to detect the substrate (upper panel) and with the anti-Strep antibody to detect Tim23 (lower panel). The substrate-crosslinked bands were detected in the upper panel, similar to the immunoblotting results in Fig. S4B. However, immunoblotting of Tim23 did not reveal any crosslinking bands in the lower panel. The non-specific bands in the immunoblots are marked by *. (**B**) The supercomplexes were assembled as in **A**, and then mitochondria were subjected to UV irradiation. Tim23 was pulled down using Ni resin under denaturing conditions. Immunoblotting with the anti-Flag antibody (upper panel) and the anti-Strep antibody (lower panel) did not detect any crosslinking bands between the substrate and Tim23. (**C**) The supercomplexes with Bpa-incorporated substrates were assembled in yeast strains expressing Strep-His tagged Tim17 (left panels, yeast strains yLS210/pLSB4-X) or Mgr2 (right panels, yeast strains yLS421/pLSB4-X). After UV irradiation of mitochondria, Tim17 or Mgr2 was pulled down using Ni resin. Immunoblotting with the anti-Flag antibody (upper panels) and the anti-Strep antibody (lower panels) detected the substrate-crosslinked band of Tim17 or Mgr2, respectively. The non-specific bands in the immunoblots are marked by *.

**Fig. S6.**
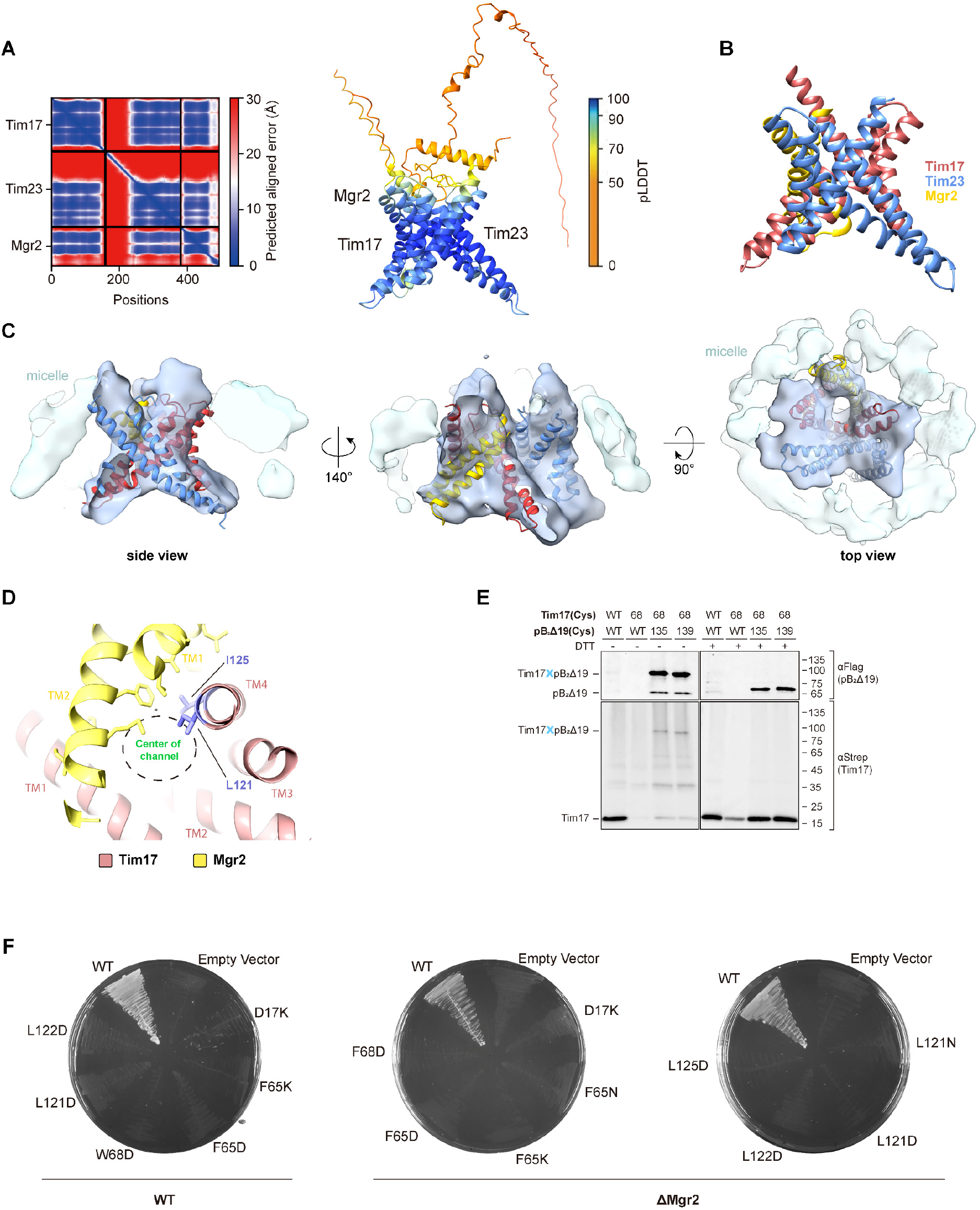
Tim23-Tim17-Mgr2 model and mutants. (**A**) AphaFold2 model of the Tim23-Tim17-Mgr2 heterotrimer, displaying the Predicted Aligned Error (PAE) plot (left) and the model (right) colored according to the predicted Local Distance Difference Test (pLDDT) score. (**B**) Model of the TM segments of Tim23-Tim17-Mgr2, showing the X shape of the heterotrimer. Tim23, Tim17, and Mgr2 are colored Dodger blue, Indian red, and yellow, respectively. (**C**) Fitting of the Tim17-Tim23-Mgr2 heterotrimer into the cryo-EM density of the TIM23 disc, shown in three views. The TIM23 density is colored sky blue and the detergent micelle density is colored light blue. (**D**) Top view of the Tim17-Mgr2 dimer, focusing on the residues 121 and 125 of Tim17, shown in three different views. Tim17 is colored Indian red and Mgr2 is colored yellow. L121 and I125 of Tim17 and the neighboring residues of Mgr2 are shown as blue and yellow sticks, respectively. The channel is indicated by a dashed circle. (**E**) Disulfide crosslinking between Tim17 and the substrate. The disulfide bond between Tim17(68C) and the substrate (135C or 139C) was detected in yeast strains yLS273/pLSB9-X. The yeast strains yLS274 and yLS276 with the wild type (WT) substrate were used as control. Tim17 was Strep-His tagged and the substrate was Flag tagged. The crosslinking products between Tim17 and the substrate were pulled down by using the Ni and Flag resins sequentially after oxidation of mitochondria by CuPh_3_. Before SDS-PAGE, the Flag eluents were split into two portions, one treated DTT and one without the DTT treatment. (**F**) Tim17 mutant selection on 5-FOA plates. The yeast strains, yLS270 and yLS280, were transformed with Tim17 mutation plasmid pLSA4-X and selected on 5-FOA plates. The mutants that could not grow during selection are shown.

**Fig. S7.**
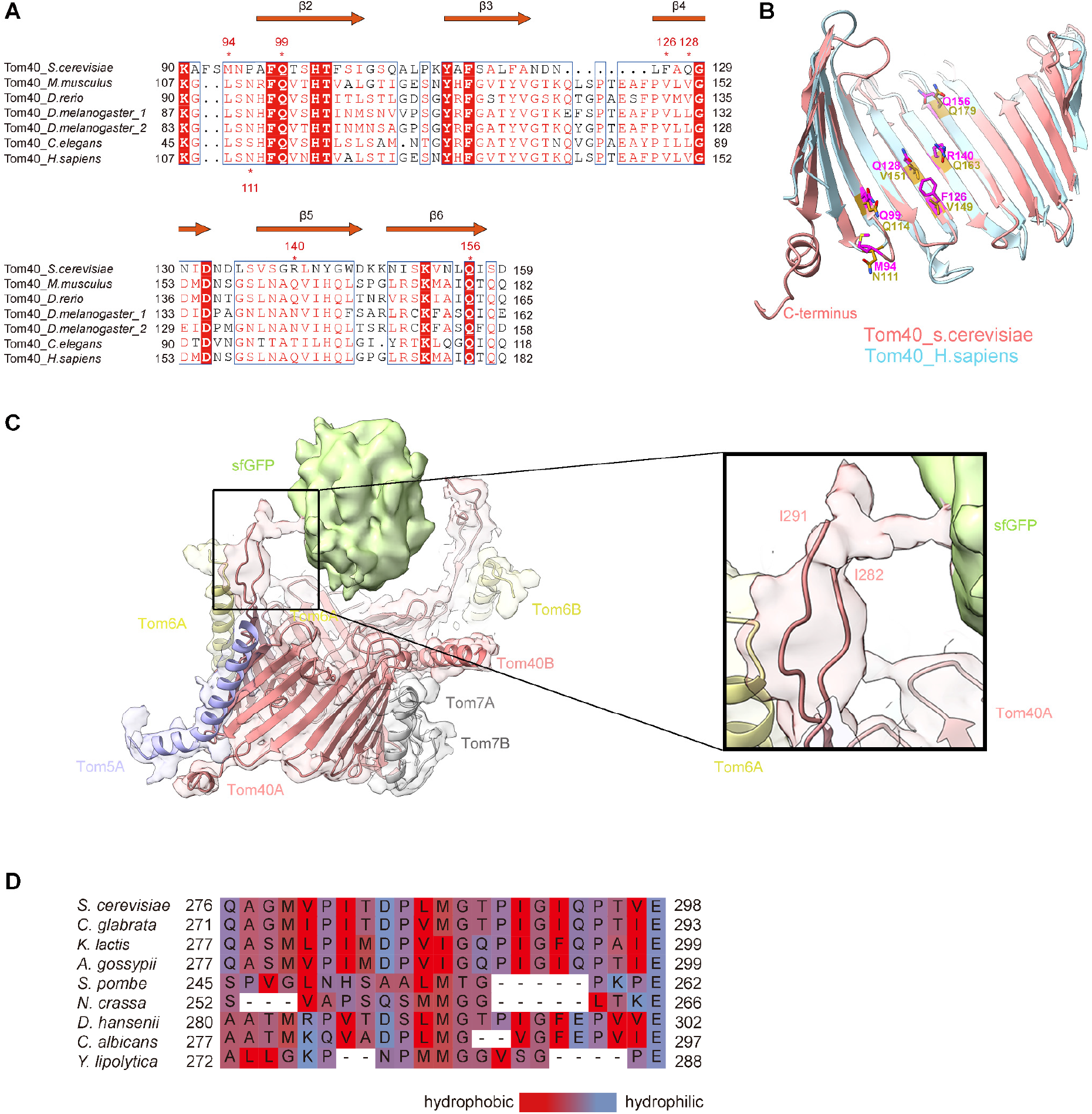
Substrate contact sites of Tom40. (**A**) Sequence alignment of Tom40 focusing on the patch2 region. The substrate interacting residues in patch2 are marked. (**B**) Structure superimposition of yeast and human Tom40. The ribbons of yeast and human Tom40 are colored salmon and cyan, respectively. The substrate interacting residues of yeast and human Tom40 are shown as sticks and colored magenta and gold, respectively. (**C**) Cryo-EM density of the substrate-engaged Tom40 channel, focusing on the contact site between GFP and L14-15 of Tom40. The tip of L14-15 (residues 283-290) is not modeled as the local density is not well resolved. However, it is clear from the map that L14-15 is in direct contact with GFP. (**D**) Sequence alignment of L14-15 from representative fungus species. The residues are colored according to their hydrophobicity.

**Fig. S8.**
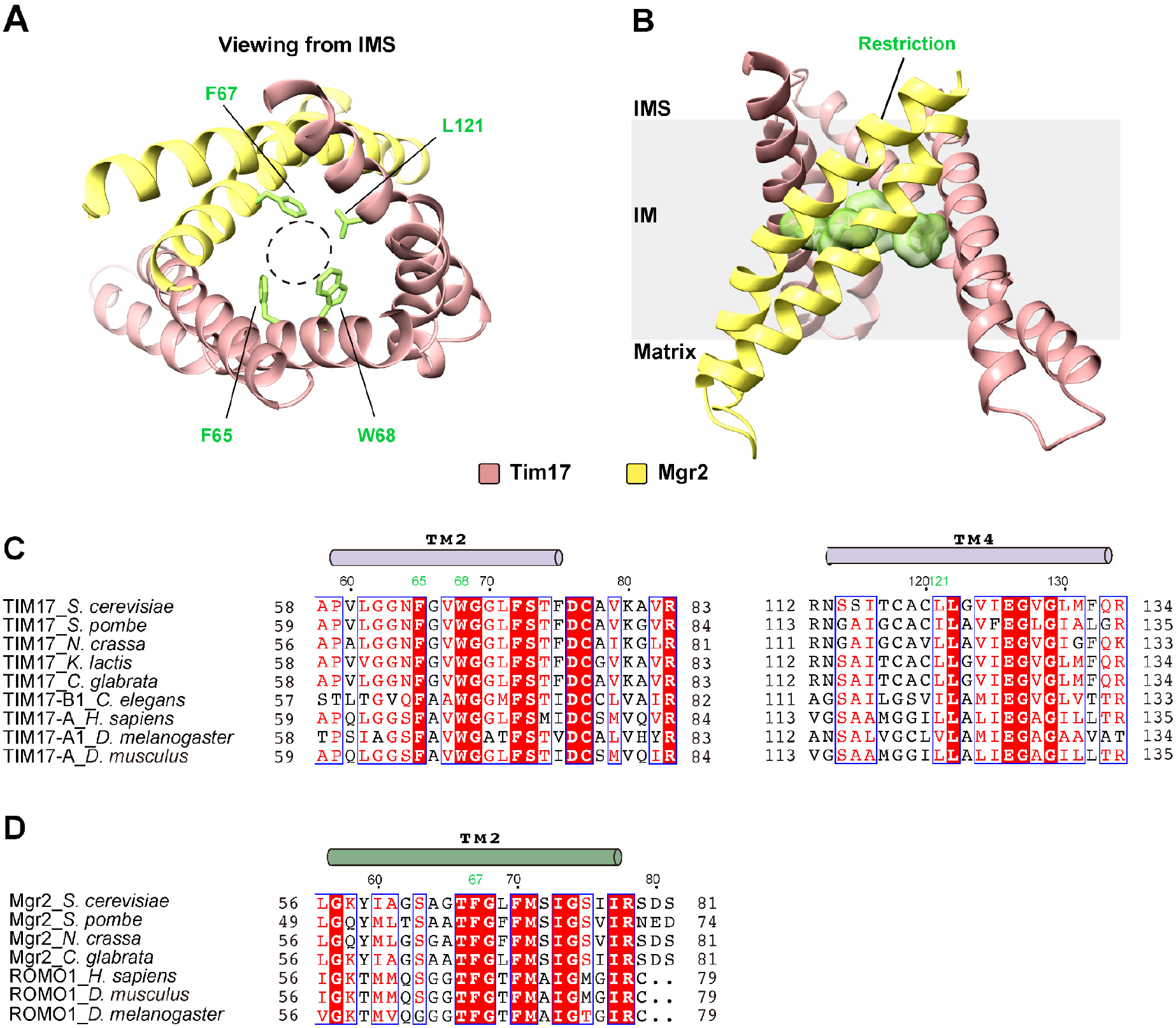
Restriction of the protein import pathway in the Tim17-Mgr2 heterodimer. (**A**) Top view of the Tim17-Mgr2 dimer, focusing on the restriction of the import pathway. The restriction residues are labeled and shown as green sticks. The opening at the restriction is indicated by a dashed circle. (**B**) Side view of the Tim17-Mgr2 dimer. The position of the restriction residues in the membrane is highlighted by green space-filling surfaces. (**C**) Sequence alignment of TM2 and TM4 of Tim17 from representative species. The purple cylinders on top of the sequences indicate the α helices of Tim17. The restriction residues are marked in green numbers. The multiple-sequence alignment was performed with ClustalOmega and visualized by ESPript. The sequences are from *Saccharomyces cerevisiae* (*S. cerevisiae,* Uniprot P39515), *Schizosaccharomyces pombe* (*S. pombe,* Uniprot P87130), *Neurospora crassa* (*N. crassa,* Uniprot P59670), *Homo sapiens* (*H. sapiens*, Uniprot Q99595), *Mus musculus* (*M. musculus*, Uniprot Q545U2), *Drosophila melanogaster* (*D. melanogaster*, Uniprot Q9VGA2), *Caenorhabditis elegans* (*C. elegans,* Uniprot O44477). (**D**) Sequence alignment of TM2 of Mgr2 from representative species. The dark green cylinders on top of the sequences indicate the α helices of Mgr2/ROMO1. The restriction residue is marked in green numbers. The sequences are from *Saccharomyces cerevisiae* (*S. cerevisiae,* Uniprot Q02889), *Schizosaccharomyces pombe* (*S. pombe,* Uniprot P79082), *Neurospora crassa* (*N. crassa,* Uniprot Q7SDL7), *Homo sapiens* (*H. sapiens*, Uniprot P60602), *Mus musculus* (*M. musculus*, Uniprot P60603), *Drosophila melanogaster* (*D. melanogaster*, Uniprot Q9VUM2), *Caenorhabditis elegans* (*C. elegans,* Uniprot Q93511).

**Fig. S9.**
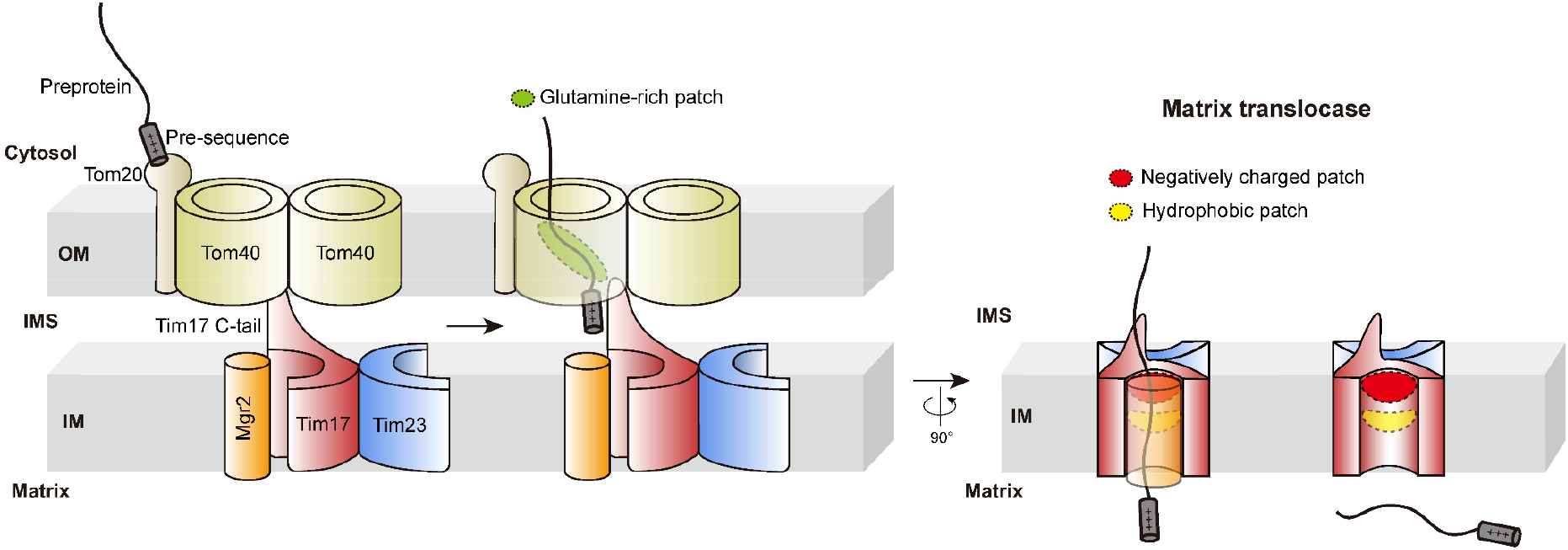
Pre-seqeunce import pathway. Scheme of mitochondrial preprotein import through the TOM-TIM23 supercomplex. The Tom40 subunit in the outer membrane, and the Tim17, Tim23, and Mgr2 subunits in the inner membrane are colored light green, Indian red, dodger blue, and orange, r espectively. The preprotein substrate is in gray. The C terminal tail of Tim17 is labeled. The glutamine-rich patch (patch 2) in Tom40 is outlined by a green oval. The negatively charged patch and hydrophobic patch in Tim17 are marked by a red and yellow oval, respectively. Mgr2 is omitted in the last panel to show the import pathway inside the concave surface of Tim17.

**Fig. S10.**
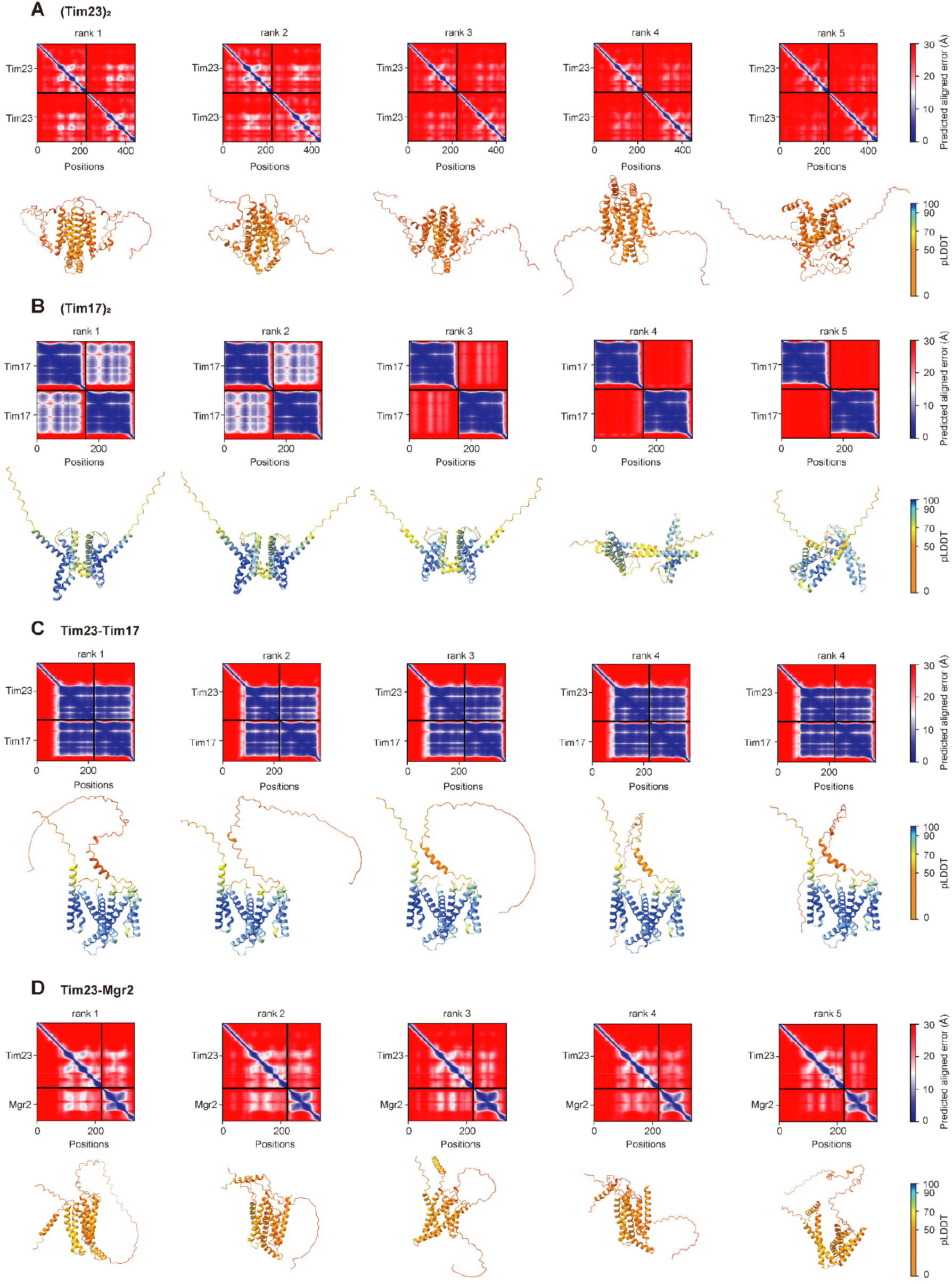
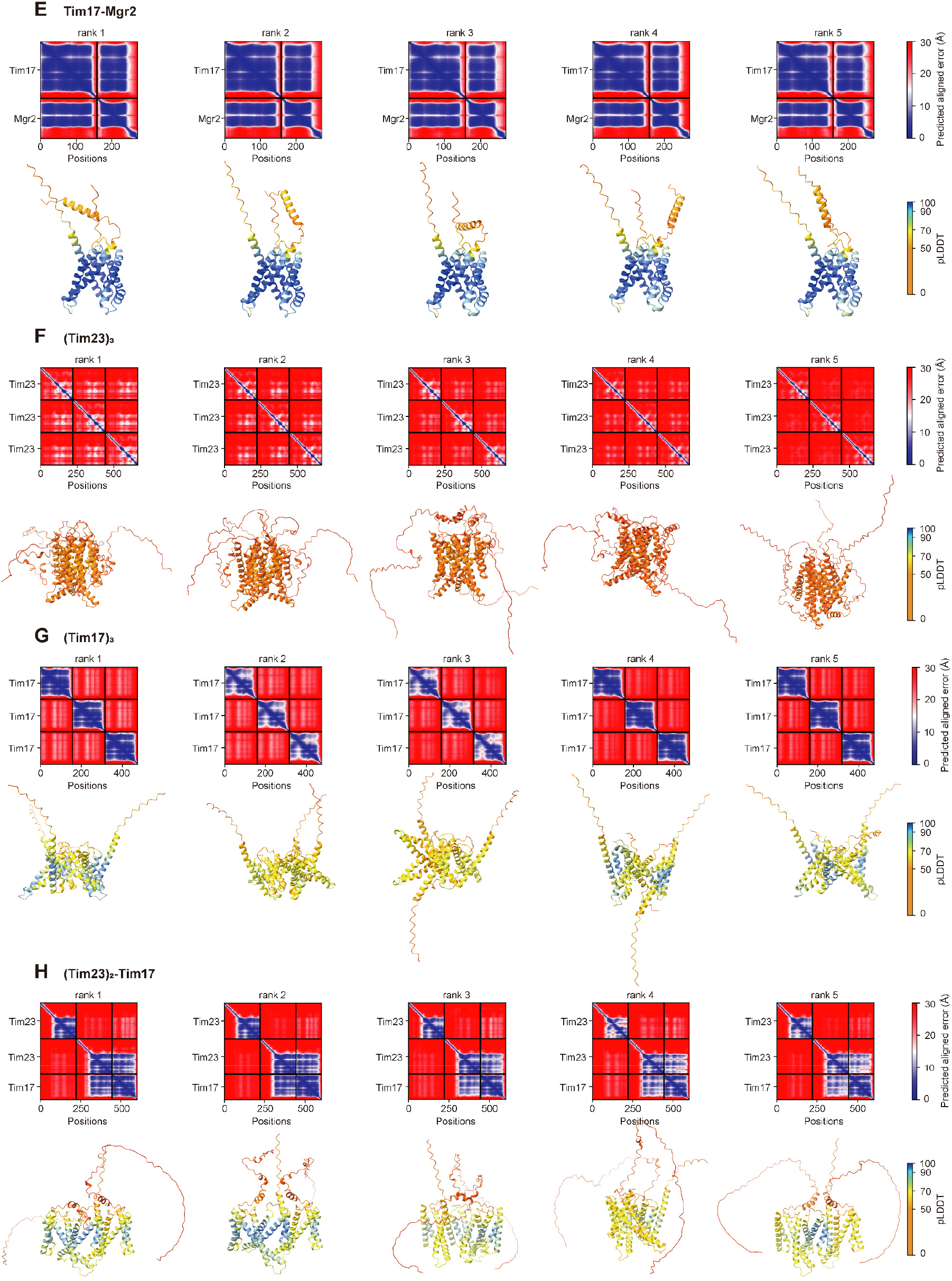
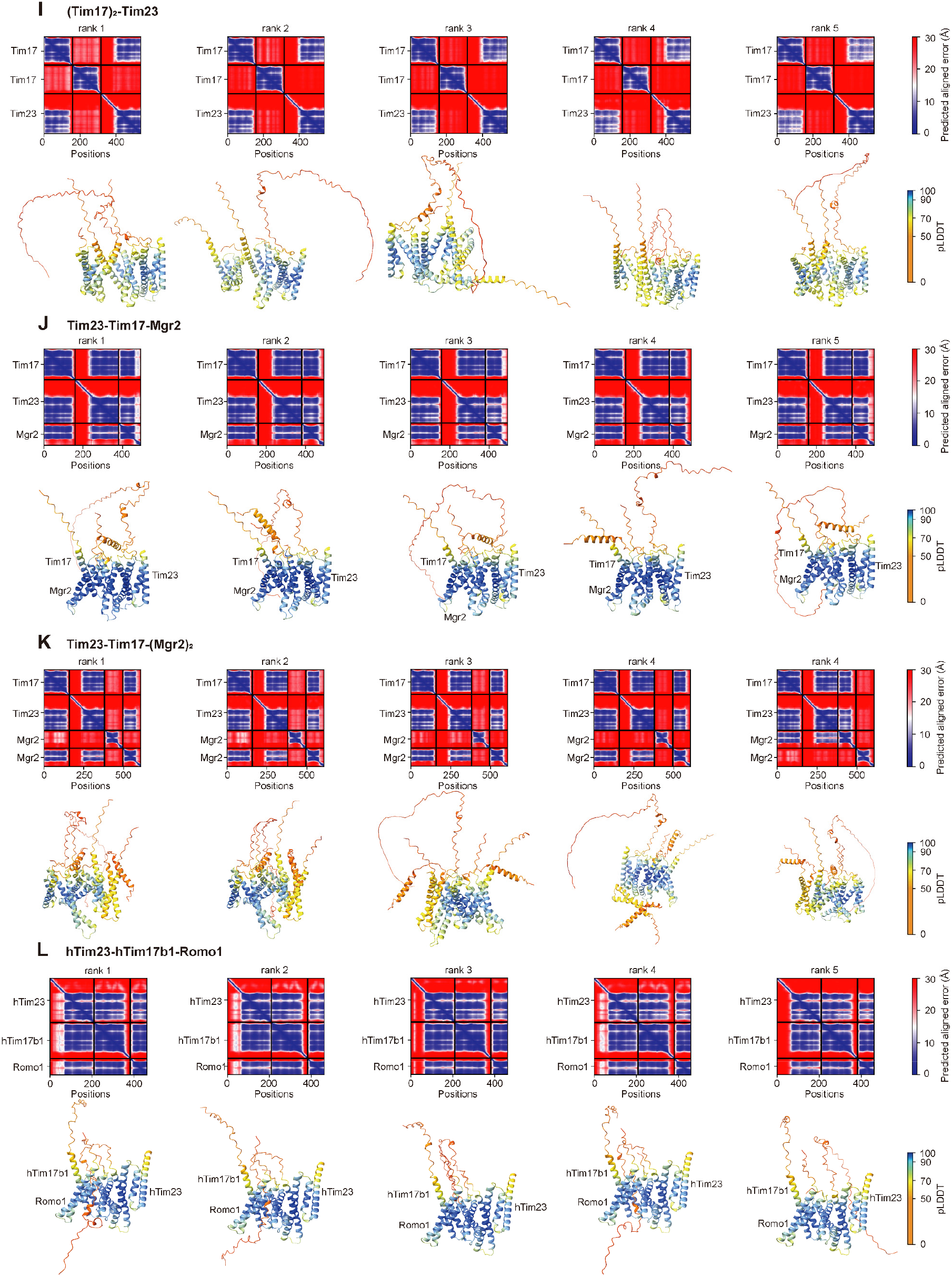
AlphaFold2 modeling of TIM23 subunits. Homo- and heter-oligomers of Tim23, Tim17, and Mgr2 are modeled by AlphaFold2. For each prediction, five models are generated and ranked on the basis of their pLDDT scores. The PAE plots and the models colored according to the pLDDT scores are displayed.

